# Monju: Multi-criteria clustering in single-cell omics

**DOI:** 10.64898/2026.05.28.728427

**Authors:** Takaaki Kaneko, Shunta Sakaguchi, Shusei Fujioka, Yuichiro Yada, Ryosuke Kojima, Honda Naoki

## Abstract

Clustering is a fundamental step in single-cell omics analysis. Although single-cell omics data can, in principle, be partitioned according to multiple biologically meaningful criteria, existing methods typically cluster cells using a single criterion. To address this problem, we developed Monju, a multi-criteria clustering method based on a deep generative mixture model. Monju divides cells into biologically reasonable submodels, each of which is equipped with an interpretable latent space. Furthermore, although the partitioning of cells into submodels varies across random seeds, each solution remains biologically plausible, collectively yielding multi-criteria clustering. Moreover, by integrating these multiple clustering solutions to perform meta-clustering, Monju enables the assessment of cluster stability. We applied Monju to human peripheral blood CITE-seq data and demonstrated that it can achieve multi-criteria clustering. Monju therefore provides a powerful and practical framework for dissecting cellular heterogeneity from multiple biological perspectives.

## 1 Introduction

In data-driven science, particularly in the field of single-cell omics, clustering is one of the most essential analytical methods[1]. Clustering, by definition, is a technique for grouping observed data according to similarity, with the goal of uncovering latent structures and patterns within the data[2]. However, there is no single criterion for how such grouping should be performed[2, 3] (**Fig. 1a**). For example, when classifying people, one could use multiple possible criteria: gender (male or female), age (child, adult, elderly), health condition (healthy, disease A, disease B), or nationality. Each of these represents a distinct way of partitioning the same population, and thus clustering can—and often should—be understood as a multi-criteria process.

**Figure 1.**
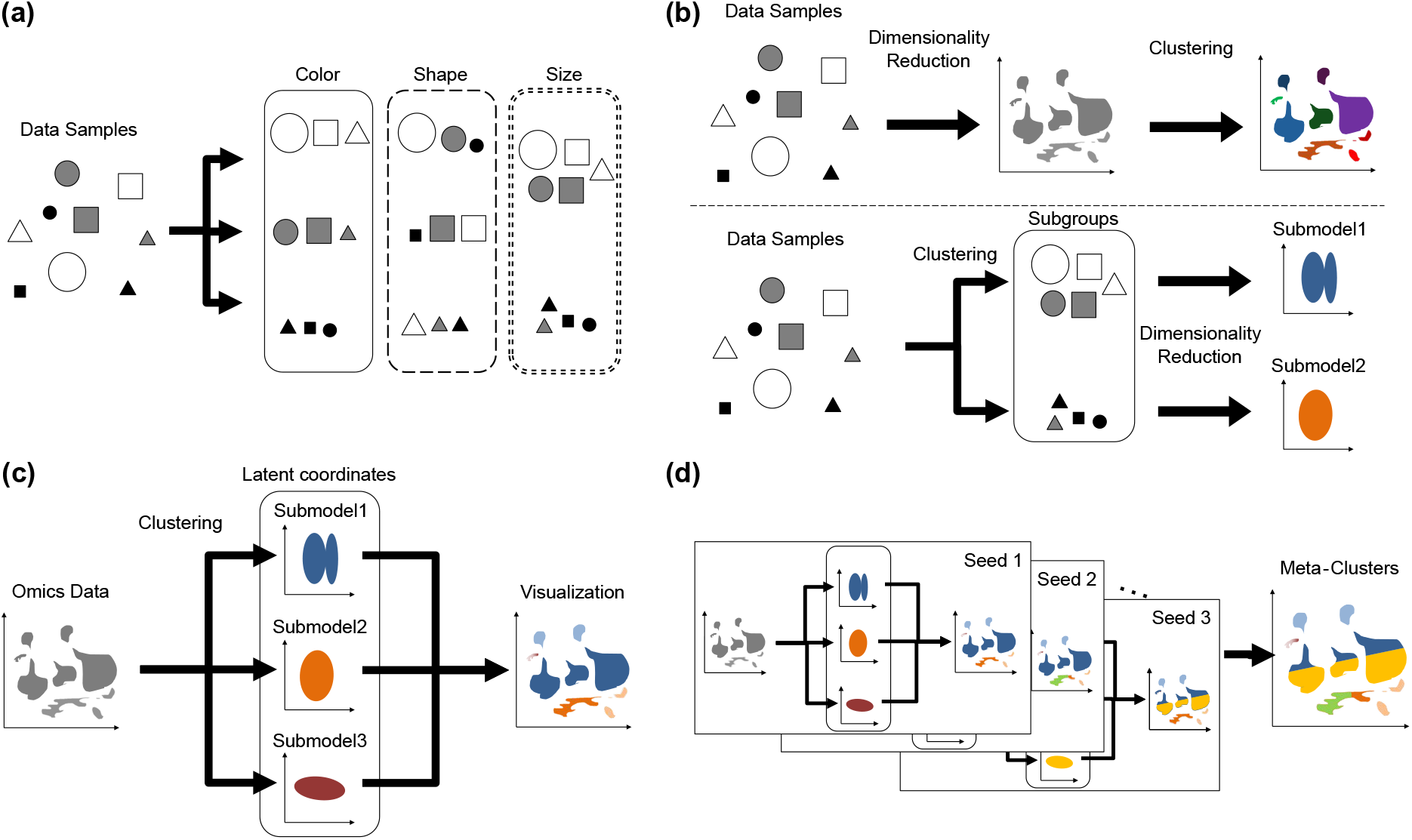
Concept of multi-criteria clustering and analytical workflow of Monju. **(a)**Schematic illustration of multi-criteria clustering. The same data can be partitioned according to different criteria, such as color, shape, or size. **(b)** Comparison of analytical strategies in single-cell omics. Top: Conventional “dimensionality reduction first, then clustering” strategy in single-cell omics analysis. Bottom: Proposed “Clustering first, then dimensionality reduction” strategy. **(c)** Overview of the Monju framework. Monju probabilistically assigns cells to multiple submodels (i.e., clustering) and embeds them into the latent space of each submodel (i.e., dimensionality reduction). Each submodel represents cells by latent coordinates that can capture biologically interpretable structures, such as functional gradients or differentiation trajectories. Visualization of submodel assignments reveals the resulting cluster structure. **(d)** The scheme of meta-clustering in Monju. Integration of clustering results across multiple training trials. Clusters obtained from independent runs with different random seeds are integrated to generate meta-clusters.

In single-cell omics, clustering plays an especially crucial role in revealing cellular heterogeneity and uncovering novel biological subpopulations[4]. However, classifying cells often requires considering multiple criteria simultaneously. Within gene expression-based analyses, cells can be grouped according to distinct biological aspects—for instance, by the expression of marker genes that define canonical cell types[5], by the activity of specific signaling pathways[6], or by profiles of metabolic gene expression[7].

Despite this complexity, most current analyses adjust clustering parameters to reproduce pre-existing definitions of cell types. Such practice, while convenient, constrains the exploratory power of data-driven approaches that are meant to minimize prior biases. Moreover, the tuning of clustering parameters is far from trivial. It is labor-intensive and typically demands substantial domain expertise[8]. To validate clustering results, researchers must annotate each cluster with putative cell-type identities, a process that depends heavily on prior biological knowledge about cell-type-specific marker genes[4, 9]. Consequently, annotation becomes a time-consuming[10] and expertise-dependent bottleneck, limiting accessibility for non-specialists and hindering large-scale, unbiased analyses.

Part of this difficulty stems not from the intrinsic properties of clustering methods themselves, but from the structure of the standard analysis pipeline[1]. Conventional approaches to single-cell omics data typically apply dimensionality reduction methods such as PCA, variational autoencoders (VAEs)[11, 12] or UMAP to gene expression data first, and then perform clustering in the resulting latent space[4, 13]. This “dimensionality reduction first, then clustering” strategy often constrains clustering to a predefined low-dimensional space, thereby limiting biological interpretability[1]. Such a framework is restricted to computing simple measures of similarity (i.e., pairwise distances) between cells after dimensionality reduction, making it difficult to classify cells according to multiple biological criteria[14, 2]. In contrast, we offer the opposite strategy—”clustering first, then dimensionality reduction.” In our strategy, cells are initially grouped into interpretable subgroups, and dimensionality reduction is then performed separately within each subgroup (**Fig. 1b**). For example, cells with high activity in a specific signaling pathway can be assigned to one subgroup, within which the reduced-dimensional axes may capture biologically meaningful variations such as differential responses of that pathway.

In this study, we developed Monju (Mixture mOdel for multi-criteria, Nonlinear, Joint Understanding) to address the challenge of multi-criteria clustering in high-dimensional omics data. Monju integrates a mixture model with deep learning to jointly infer cluster assignments and latent coordinates within each submodel (**Fig. 1c-d**). By decomposing heterogeneous cell populations into functionally interpretable submodels (i.e., “clustering first”) and performing dimensionality reduction within each context (i.e., “then dimensionality reduction”), Monju can identify structures that extend beyond conventional cluster boundaries. Using human peripheral blood mononuclear cells (PBMC) CITE-seq data, we show that Monju recovers biologically meaningful partitions and interpretable submodel-specific latent structure while also providing a framework to examine clustering variability across runs and cellular heterogeneity in the proposed Monju Space.

## 2 Results

### 2.1 Monju model

Monju is a deep generative mixture model. We assume that single-cell omics data consist of multiple subpopulations, each of which can be described by its own low-dimensional latent structure. Under this assumption, understanding the data requires estimating both (i) which subpopulation each cell belongs to (i.e., clustering) and (ii) where the cell is located within the latent space of that subpopulation (i.e., dimensionality reduction).

Monju takes raw gene-expression count data as input. A neural network module partitions cells into multiple submodels, corresponding to clustering, while each submodel uses a variational autoencoder (VAE) to learn a low-dimensional latent space for the cells assigned to it. Thus, Monju simultaneously performs clustering and submodel-specific dimensionality reduction (**Fig. 3a**). The partitioning neural network and all submodel-specific VAEs are trained jointly in an end-to-end manner so that the entire dataset is well reconstructed. Furthermore, to handle multimodal data, such as RNA and protein measurements, Monju adopts a mixture-of-experts (MoE) architecture (see Methods: Monju model formulation).

**Figure 2.**
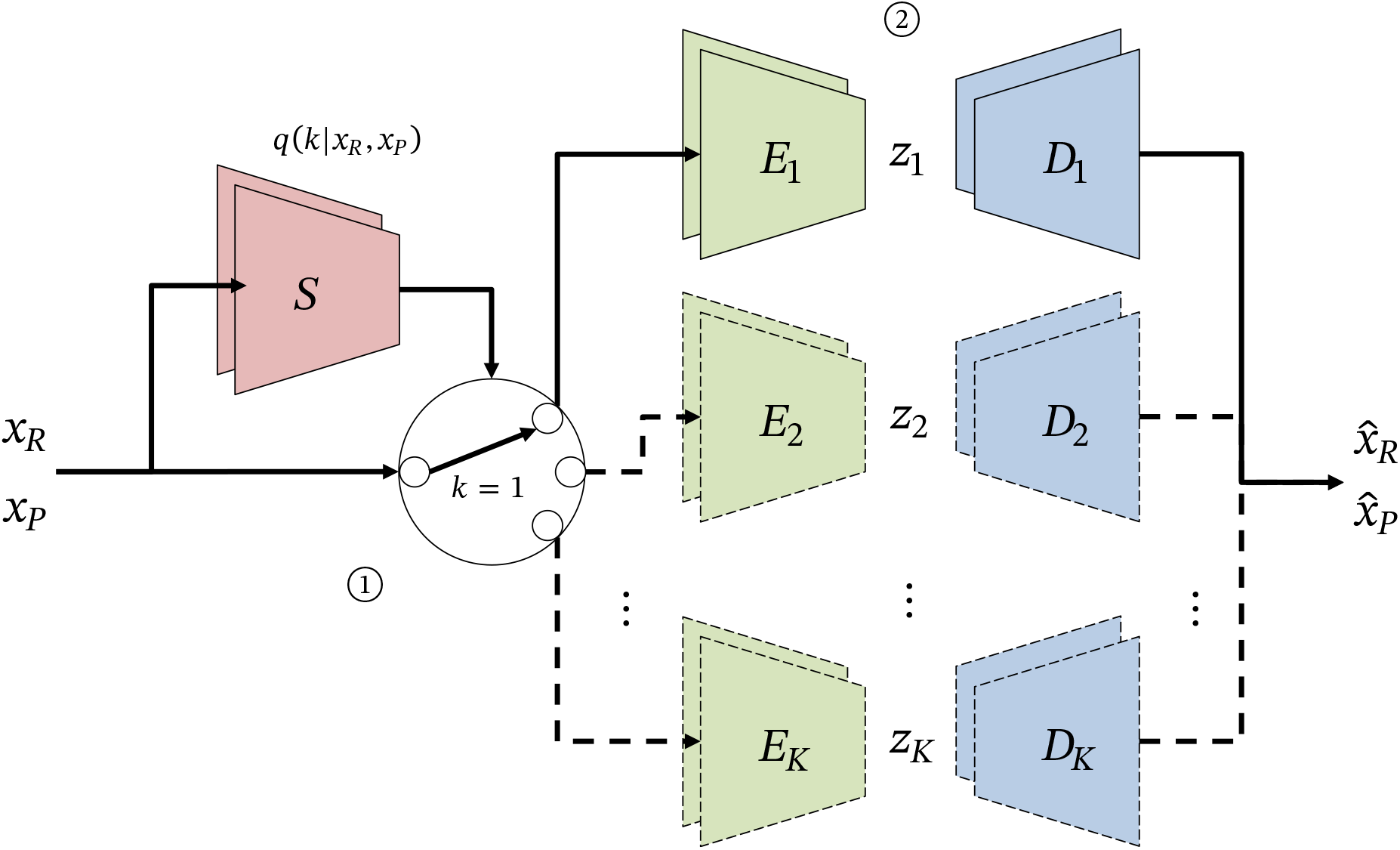
Monju architecture. **(1)** A classifier network *S* takes multimodal input data (*x*_*R*_, *x*_*P*_ ) and outputs a probabilistic assignment *q*(*k*| *x*_*R*_, *x*_*P*_ ) over *K* submodels. Each cell is stochastically assigned to one of the submodels. **(2)** Each submodel consists of a VAE composed of an encoder *E*_*k*_ and a decoder *D*_*k*_, which maps the input data to a low-dimensional latent representation *z*_*k*_ and reconstructs the input as 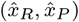, respectively. The model is trained end-to-end to maximize reconstruction accuracy across all submodels. Through this process, the classifier learns to assign cells to the submodel that best reconstructs them, effectively grouping cells that share a common low-dimensional generative structure.

**Figure 3.**
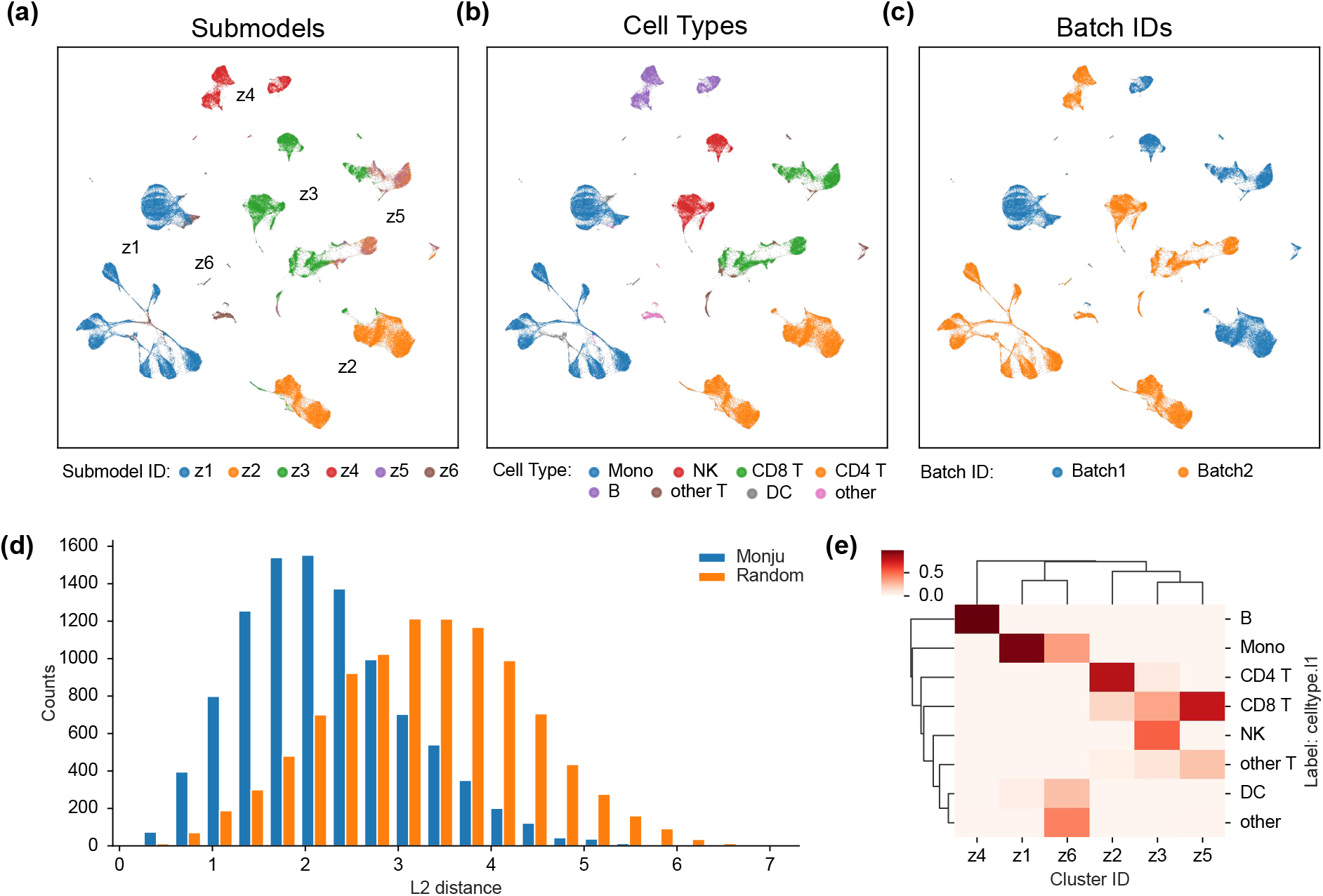
Monju assigns cells to biologically meaningful submodels. **(a,b, and c)** UMAP visualization for PBMC CITE-seq data based on the latent space of a conventional non-mixture VAE method. Cells are colored by Monju submodel assignment **(a)**, annotated cell type labels (celltype.l1; **(b)**), and batch ID **(c).** Although Monju was trained with *K* = 20 submodels, only submodels with assigned cells are shown in the figure. Monju assigned spatially adjacent cell populations to the same submodels, and these assignments closely resembled known cell-type distributions rather than batch structure. **(d)** Distribution of pairwise distances in the conventional VAE latent space between cells assigned to the same Monju submodel and between randomly selected cells. Cells sharing the same Monju submodel showed shorter distances than randomly paired cells, indicating that Monju groups nearby cells in the reference latent space. **(e)** Cell-type composition of each Monju submodel. Most submodels were predominantly composed of a single annotated cell type, supporting the biological interpretability of the assignments. Notably, submodel *z*_3_ contained both NK cells and a subset of CD8^+^ T cells, suggesting that Monju can capture functional similarity, such as cytotoxicity, beyond conventional cell-type boundaries.

Importantly, because each submodel is implemented as a VAE with a relatively simple latent representation, the model is encouraged to assign each cell to the submodel that can reconstruct it most effectively. In other words, the classifier learns to assign cells that are well described by a common low-dimensional structure to the same VAE, thereby improving reconstruction efficiency. This mechanism defines clustering based not on pairwise similarity in a fixed space, but on how well cells can be explained by shared generative structure.

Because the partitioning pattern is not unique in general (as seen in **Fig. 1a**), Monju learns different partitions and submodel-specific latent spaces across trials with different random seeds. Thus, each trial yields a distinct clustering criterion. By repeating the process, Monju generates multiple clustering results based on different criteria. These results can then be integrated through meta-clustering to obtain a comprehensive representation that incorporates multiple partitioning criteria.

### 2.2 Monju divided cells into biologically reasonable submodels (Fig. 3)

We applied Monju to single-cell RNA and surface-protein expression data obtained from the PBMC CITE-seq dataset[15] (see Methods: Dataset). As a first step, we verified that Monju, as a mixture model, can perform proper learning comparable to a non-mixture, conventional VAE[12]. We confirmed that Monju successfully achieved reconstruction and cross-modal prediction within each submodel (**Fig. S1**). Next, we examined whether Monju reasonably assigned cells to each submodel. We analyzed the assignment weights of each cell across submodels and found that the maximum assignment weight was close to 1 for nearly all cells (**Fig. S2**), indicating an almost one-hot assignment pattern. This result suggests that Monju assigns each cell predominantly to a single submodel.

To assess whether the submodel assignments captured meaningful structure, we visualized Monju submodel assignments on a UMAP plot based on the latent space provided by a conventional non-mixture VAE method[12] (**Fig. 3a**). We observed that spatially adjacent cell populations in UMAP space based on the conventional non-mixture VAE method were assigned to the same submodel. To quantitatively validate this, we calculated pairwise distances in the latent space of the non-mixture VAE between cells assigned to the same submodel. Comparing their distribution with that obtained from randomly assigned cells (see Methods and **Fig. 3** for the plotting procedure), we showed that cells sharing the same submodel showed markedly smaller distances (**Fig. 3d**).

We then compared the submodel assignments with annotated cell types. When cell-type labels (celltype.l1) were overlaid on the same UMAP (**Fig. 3b**), their distribution closely resembled that of the submodels. The composition of cell types within each submodel confirmed that most submodels consisted predominantly of a single cell type (**Fig. 3e**). In contrast, visualization by batch ID (**Fig. 3c**) showed that most submodels contained cells from both batches, indicating that Monju’s assignments were driven by biological similarity rather than batch effects. These results indicate that Monju’s submodel assignments correspond well to known cell types. Interestingly, one of the submodels *z*_3_ included both NK and a subset of CD8^+^ T cells, while other subsets of CD8^+^ T cells were contained in other submodels (**Fig. 3e**). Since NK and CD8^+^ T cells share strong cytotoxic functions and exhibit similar gene- and protein-expression profiles[16, 17], Monju can capture a functional substructure rather than strict cell-type boundaries. Taken together, these findings demonstrate that Monju performs biologically meaningful decomposition of cellular populations. Additional comparisons with other data labels are provided in Supplementary **Figs. S3** **and S4**.

### 2.3 Monju extracted biologically interpretable coordinates in submodels (Fig. 4)

We visualized the distribution of cells assigned to each submodel to assess their biological interpretability (**Fig. 4 and Supplementary Figs. S5 to S15**). Focusing on the submodel *z*_2_, which mainly contained CD4^+^ T cells, we found that the cells were linearly arranged in the PCA space (arrow in **Fig. 4a**), rather than forming a complex manifold as in the original UMAP space. We termed the axis indicated by this arrow the mixed principal component (mPC) axis (see Methods: Analysis of submodel assignments). Cell labels revealed an ordered continuum from naïve T cells to CD4^+^ central memory T (TCM) cells and CD4^+^ effector memory T (TEM) cells, suggesting that the mPC axis reflects a biologically functional gradient of T cells. Moreover, an axis orthogonal to this mPC corresponded to batch differences (**Fig. 4b-c**), indicating that the submodel represents batch effects as an axis independent of biological function.

**Figure 4.**
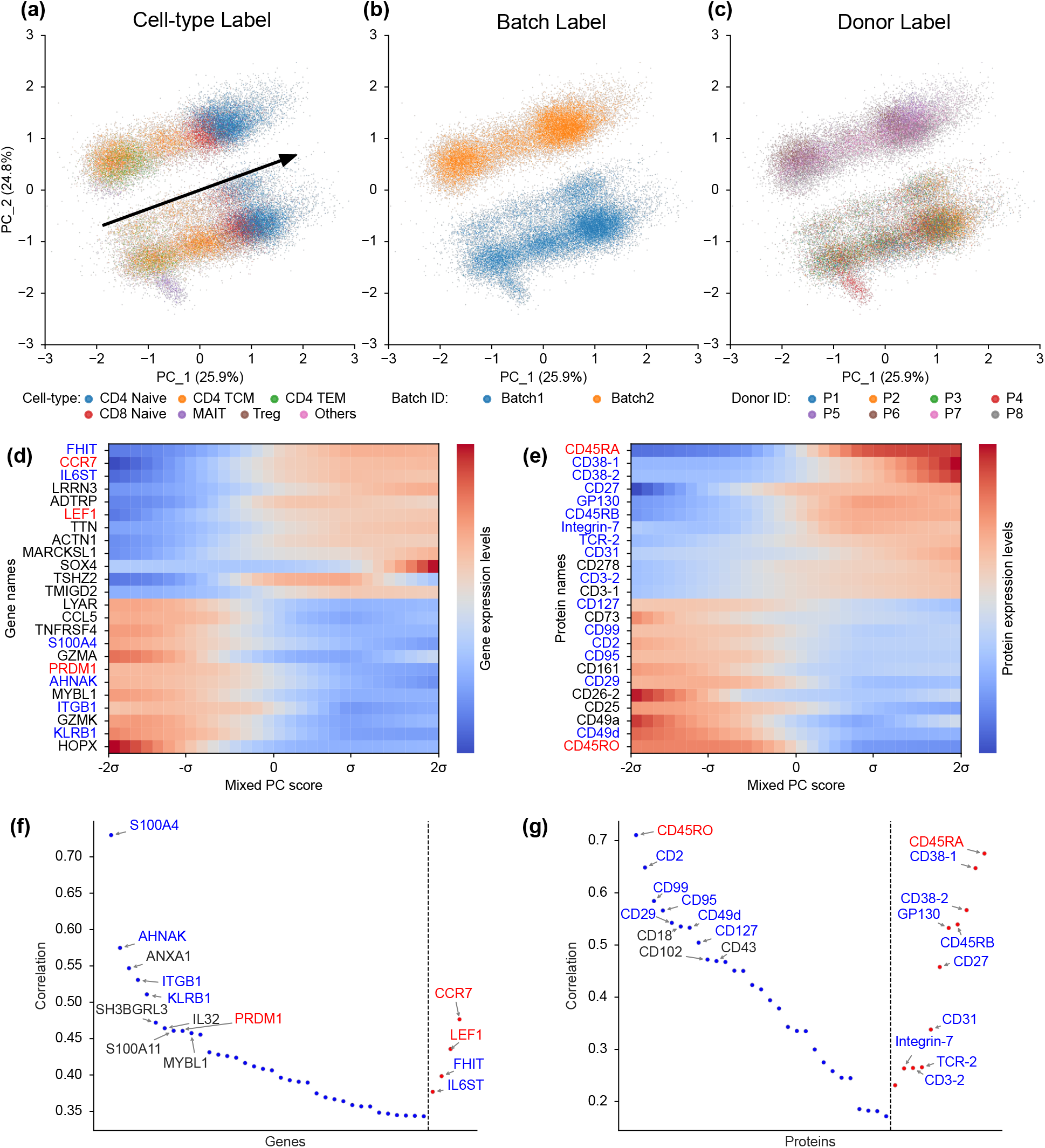
Interpretability of the latent representation learned by Monju. **(a,b, and c)** PCA visualization of submodel *z*_2_. Only cells assigned to *z*_2_ were extracted, and PCA was applied to their latent representations within this submodel. Points represent individual cells and are colored by their **(a)** cell-type labels, batch ID, and **(c)** donor ID, respectively. The arrow in (a) shows the axis used for visualizing expression changes in panel (d and e). **(d and e)** Heatmaps showing reconstructed gene **(d)** and protein **(e)** expression along the axis indicated in (a), as generated by the submodel decoder. The horizontal axis represents positions along the axis. Expression values at each position were estimated using the corresponding submodel VAE by decoding latent coordinates sampled along this axis. The vertical axis lists the top 12 genes (or proteins) with the largest positive covariance and the top 12 with the largest negative covariance between reconstructed expression and mPC position, ordered from negative to positive (bottom to top). Labels shown in blue indicate molecules that also appear in (f and g), whereas labels shown in red indicate molecules discussed in the main text. Expression values are shown on a log scale and normalized to have a mean of zero for each gene or protein. **(f and g)** Genes **(f)** and proteins **(g)** whose observed (i.e., measured, not model-reconstructed) expression levels correlate with the mixed principal component in (a). Genes and proteins are ordered along the horizontal axis by ascending Spearman correlation coefficient. The vertical axis indicates the absolute Spearman correlation between the mPC score and observed expression. Positive correlations are shown in red and negative correlations in blue. Only the top 40 genes and top 40 proteins with the largest absolute correlations are displayed.

To examine whether the mPC axis corresponds to molecular changes associated with T-cell functional states, we performed two complementary analyses. We first analyzed how gene and protein expression are generated by Monju along the mPC axis. We ranked genes and proteins by the covariance between reconstructed expression and mPC position and visualized those with the largest values (**Fig. 4d-e**). For gene expression, CCR7 and LEF1, which are known to be highly expressed in naïve and central memory T cells[18, 19, 20], increased toward the positive end of the mPC axis. In contrast, PRDM1 (Blimp-1), which drives effector differentiation of CD4^+^ T cells[21, 22], increased toward the negative end. This pattern indicates that the mPC axis captures a graded shift in gene expression corresponding to T-cell functional states. For surface proteins, CD45RA increased in the positive direction, whereas CD45RO increased in the negative direction. Because naïve T cells are CD45RA^+^CD45RO^*−*^, and TCM/TEM cells are CD45RA^*−*^CD45RO^+^[19, 20], this result mirrors the gene-expression trend and suggests that the mPC axis also reflects protein-level changes associated with T-cell functional states.

We next asked whether the same molecular gradient is directly detectable in the raw measurements, without invoking the generative model. As a standard non-parametric criterion, we computed Spearman rank correlations between the mPC score and observed expression levels, and visualized the top 40 genes and top 40 proteins ranked by |*ρ*| among features with a false discovery rate (FDR) below 0.05 (**Fig. 4f-g**). Most of these molecules overlapped with those identified in **Fig. 4d-e** (labels highlighted in blue or red in panels d and e), indicating that the gradient captured by Monju is present in the observed data and is not an artifact of the generative model. Together, these findings indicate that the submodels learned by Monju possess high interpretability, capturing coherent molecular gradients that correspond to known biological functions.

### 2.4 Assessing Cluster Stability by Meta-clustering (Fig. 5)

Up to this point, we have discussed clustering based on a single criterion. However, such a result represents only one of many possible clusterings of cells, and how cells are divided is not uniquely defined in general; multiple clustering criteria can therefore coexist (**Fig. 1a**). To accommodate this non-uniqueness, Monju leverages the inherent stochasticity of deep learning models, which leads to different outcomes across runs, thereby generating multiple criteria for partitioning the data. We confirmed that Monju runs with different random seeds produced diverse submodel partitions. Importantly, these clustering patterns corresponded to biologically meaningful distinctions in terms of cell types and functional states (**Fig. S16**).

**Figure 5.**
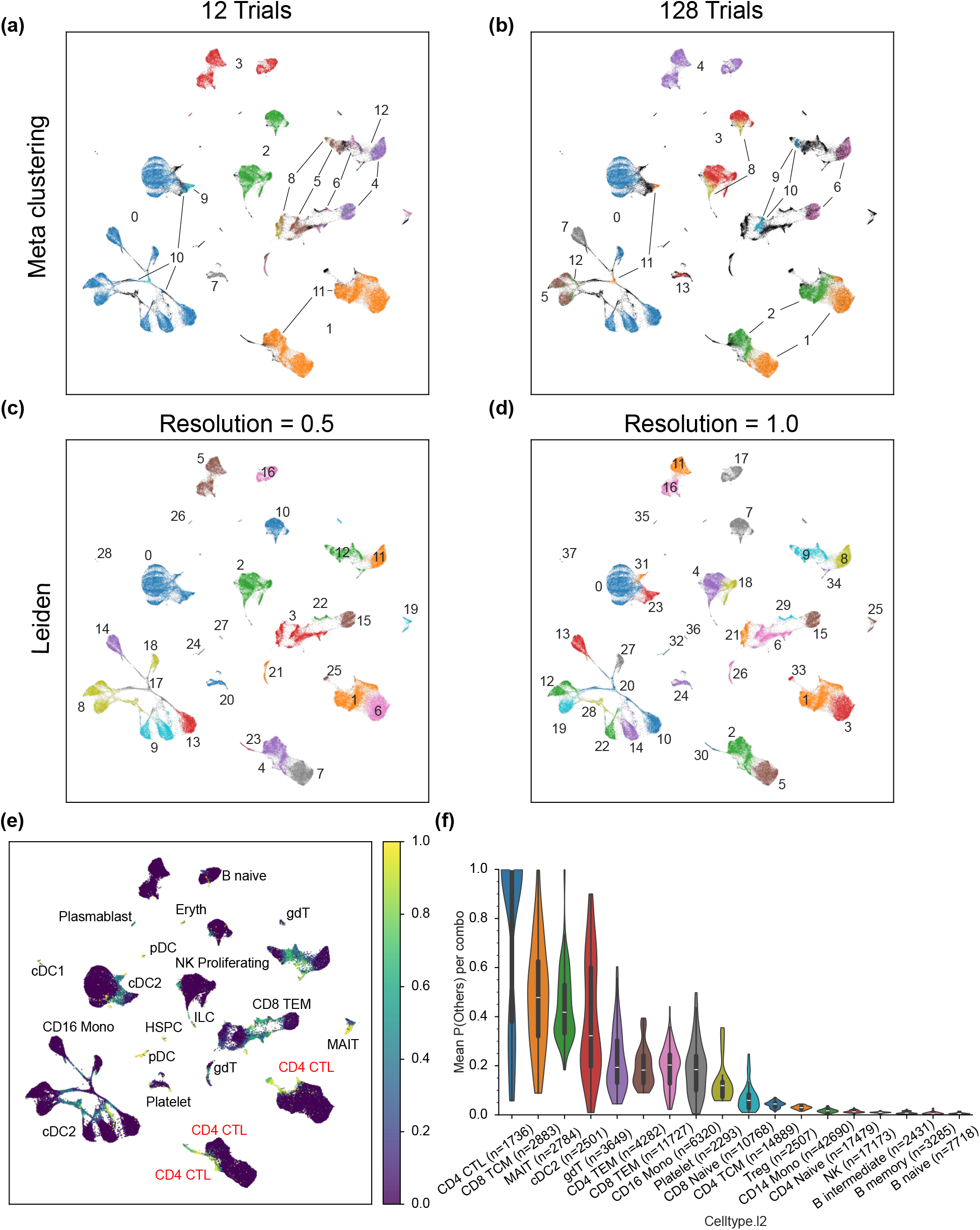
Evaluation of meta-cluster stability. **(a and b)** Monju meta-clusters (with 12 and 128 trials). The same latent space as in Fig. 3a is visualized by UMAP. Colors represent the assigned Monju meta-cluster ID. Meta-clusters containing fewer than 1,000 cells are collectively labeled as Others. **(c and d)** Leiden clusters in the same latent space (resolution parameter = 0.5 and 1.0). Colors represent the assigned Leiden cluster ID. **(e)** Quantification of meta-clustering ambiguity. The frequency of being assigned to Others in Monju meta-clustering (with 12 trials) was visualized by a color map. **(f)** Violin plots of the frequency of being assigned to Others for each cell type (celltype.l2).

Across trials, some cell populations showed high consistency in their submodel assignments. In particular, CD4^+^ T cells, NK cells, and B cells were consistently assigned to the same sub-models, respectively, regardless of the random seed, indicating that these cell types form robust populations across multiple clustering criteria. In contrast, CD8^+^ T cells exhibited substantial variability across trials. In some runs, CD8^+^ T cells formed an independent submodel (e.g., Trial 5), whereas in other runs they were grouped together with CD4^+^ T cells as a general T-cell population (e.g., Trial 8). In additional trials, all CD8^+^ T cells were grouped with NK cells as cytotoxic cells (e.g., Trial 7). Furthermore, as discussed above, only effector CD8^+^ T cells, which have high cytotoxicity, were grouped together with NK cells, while other CD8^+^ T cells remained separate (e.g., Trial 2). Thus, CD8^+^ T cells did not form a single consistent submodel across trials.

We hypothesized that integrating these clustering results would enable clustering that reflects multiple criteria, and therefore performed meta-clustering using the following procedure. In a single Monju run, each cell is assigned a submodel label. By repeating the analysis multiple times, each cell acquires a sequence of submodel labels corresponding to the number of trials. We then performed meta-clustering, defined here as clustering over clustering results, by grouping cells that shared identical label sequences across trials and treating them as belonging to the same cluster (see Methods: Monju model formulation).

As the number of runs increased, Monju meta-clustering exhibited a distinct tendency in which very small meta-clusters (fewer than 1,000 cells; “Others” in **Fig. 5a-b**) gradually separated from large and well-established clusters. These small meta-clusters arise because some cells are grouped with different cell populations depending on the criterion used in each run. This behavior is characteristic of meta-clustering. By contrast, in conventional clustering methods such as the Leiden algorithm[23], tuning the resolution parameter toward finer partitions typically resulted in the data being divided into clusters of comparable sizes (**Fig. 5c-d**).

We reasoned that this property could be used to examine the stability and variability of cell-state organization across multiple biological criteria, thereby providing insight into both well-defined cell-type boundaries and continuous functional transitions. To test this idea, we quantified meta-clustering ambiguity as follows. We first ran Monju with 128 different random seeds, generating 128 independent clustering results. We then repeatedly constructed meta-clusters by randomly selecting 12 distinct trials without replacement and integrating their results. This random selection was repeated 100 times. For each cell, we computed how frequently it was assigned to the Others category across these 100 meta-clustering runs (**Fig. 5e-f** ).

The frequency with which cells were assigned to the **Others** category varied substantially across cells. Notably, cells with a high tendency to be classified as Others were enriched at the boundaries between naïve, central memory (TCM), and effector memory (TEM) states within CD8^+^ T cells, as well as at the boundary between CD14^+^ and CD16^+^ monocytes[24]. This pattern suggests that transitions between these states are not sharply delineated but instead occur in a continuous manner. In contrast, B cells were rarely assigned to the Others category. This pattern may suggest that this cell population represents a cell type that is more discretely separated from the other clusters. In addition, CD4^+^ cytotoxic T lymphocytes (CTLs) also exhibited a high assignment frequency to **Others**. Although these cells are classified as CD4^+^ T cells, they possess cytotoxic features and display intermediate characteristics between CD4^+^ and CD8^+^ T cells[25]. This observation is consistent with the interpretation that the assignment frequency to Others serves as an indicator of state ambiguity. Taken together, these results suggest that seed-to-seed variability in Monju, when aggregated through meta-clustering, can be used not merely as a source of stochastic noise but as an informative signal for quantifying the discreteness of cell populations and the continuity or ambiguity of cellular states.

### 2.5 Monju Space: Global visualization of cell populations (Fig. 6)

In the analyses above, visualization was performed separately within the low-dimensional spaces learned by individual submodels or by a conventional VAE. While this approach enabled detailed interpretation within each submodel, it did not provide a global view of all cells across submodels. To address this limitation, we focused on the classifier network in Monju (*S* in Fig. 2), specifically the assignment probability vector produced for each cell. For each cell, we concatenated the assignment probability vectors obtained across multiple runs with different random seeds into a single vector representation (**Fig. 6a**). We defined the resulting representation space as Monju Space.

**Figure 6.**
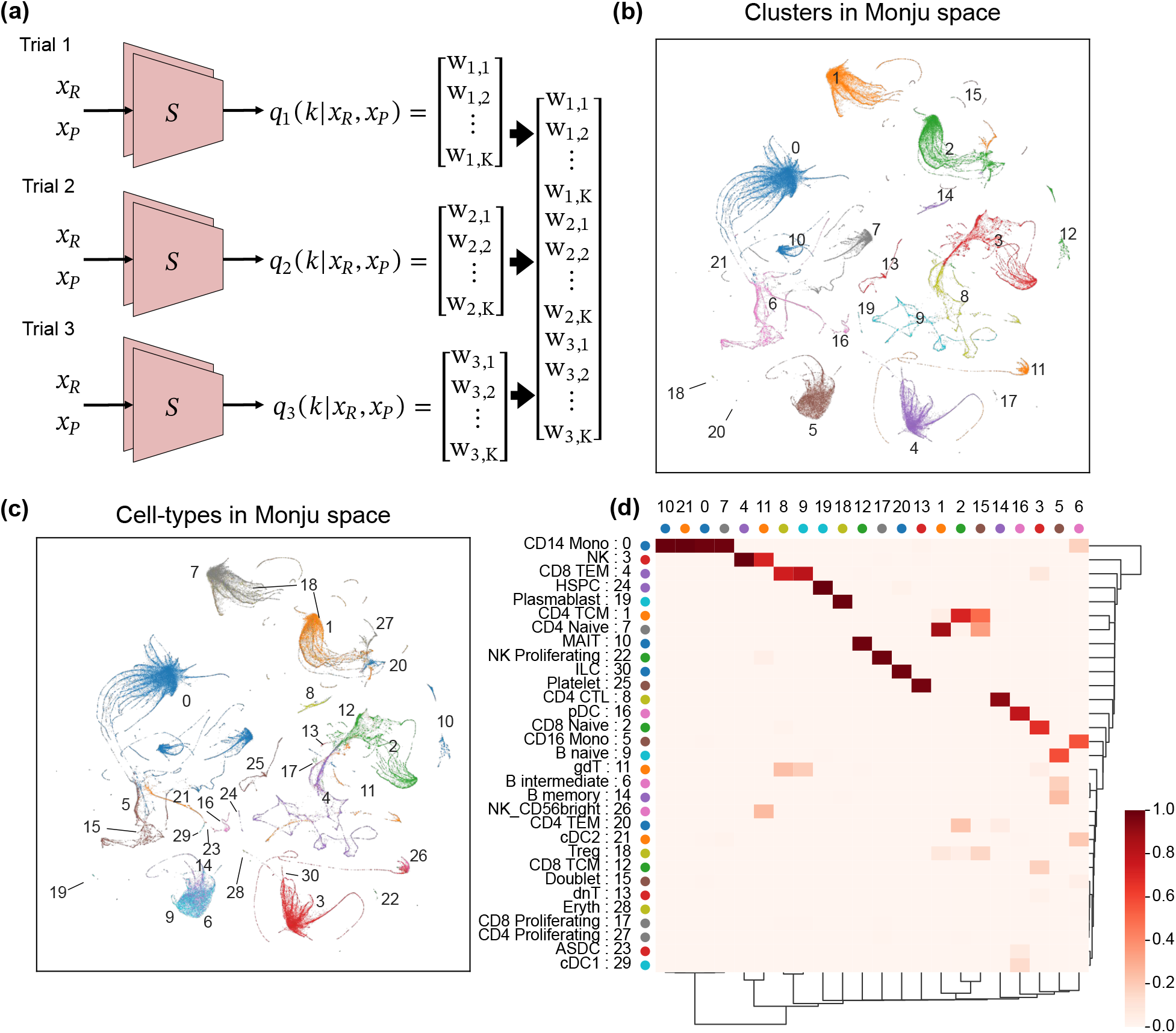
Visualization of the Monju Space. **(a)** Workflow for constructing and visualizing the Monju Space. For each cell, the assignment probability vector from classifier network *S* obtained from each trial is concatenated across all trials to form a single vector in the Monju Space. **(b and c)** The Monju Space visualization by UMAP. Colors of dots represent Leiden clustering (b) and cell-type (celltype.l2) labels (c), respectively. **(d)** Proportions of cell-type labels (celltype.l2) within each Leiden cluster in the Monju Space. The vertical axis corresponds to the legend in (c), and the horizontal axis corresponds to the legend in (b).

To assess whether the Monju Space provides a meaningful representation, we visualized it using UMAP and colored cells by Leiden cluster labels and annotated cell-type labels (**Fig. 6b,c**). The cell-type labels showed that distinct cell types were well separated in Monju Space. The Leiden clusters were highly concordant with existing cell-type annotations, as shown by their cell-type composition (**Fig. 6d**). Taken together, these results indicate that Monju Space appropriately captures cellular heterogeneity.

## 3 Discussion

In this study, we developed Monju, a deep generative mixture model that enables data-driven discovery of biologically meaningful submodels from single-cell omics data without requiring prior biological annotations. Using PBMC CITE-seq data[15], we showed that Monju identifies submodels that reflect functional relationships among cells, such as cytotoxic similarity between NK cells and CD8^+^ T cells. In addition, latent axes within each submodel were biologically interpretable, suggesting that the submodels captured meaningful biological structure rather than merely improving predictive fit. We further showed that Monju exhibits run-to-run variation in submodel partitioning, and that this property can be leveraged to evaluate cluster stability and the continuity of cellular states. Finally, we introduced the Monju Space, a partition-driven representation constructed by integrating assignment probability vectors across multiple Monju runs, and demonstrated that it appropriately captures cellular heterogeneity.

A key motivation for this work is that single-cell omics data should be partitioned according to multiple biologically meaningful criteria[2, 3]. In practice, however, most analyses rely on a single dominant clustering objective, typically corresponding to separating canonical cell types[5]. As a result, alternative biological structures—such as functional states or differentiation continua—may be overlooked or forced to conform to predefined cell-type boundaries. Moreover, the interpretation of clustering results typically depends on manual annotation using known marker genes, which requires substantial domain expertise and limits accessibility for non-specialists. These limitations highlight the need for approaches that can discover and integrate multiple clustering criteria in a data-driven manner.

A key feature of Monju is that it jointly infers cell partitioning and submodel-specific latent representations. This contrasts with the standard workflow in single-cell omics analysis, in which dimensionality reduction is performed first and clustering is applied afterward. In such workflows, once cells are embedded into a reduced space, downstream clustering is constrained by the distances or neighborhoods defined in that space. This can be limiting when multiple notions of similarity are biologically relevant, such as cell identity, functional state, and signaling activity. By learning cell partitioning and submodel-specific latent spaces simultaneously, Monju avoids reliance on a single global embedding and instead captures multiple biologically meaningful structures within the same dataset.

This joint design also facilitates interpretation at the submodel level. Because cells are modeled within relatively homogeneous subsets, the learned latent axes can be more directly related to biological variation. For example, Monju automatically identified an functional-state axis within CD4^+^ T cells. In a conventional pipeline, extracting such an axis would typically require multiple manual steps, including clustering, annotation, knowledge-based subset selection, and re-embedding[4]. Monju provides a more data-driven and automated alternative by learning subgroup-specific latent structures as part of the model itself.

Moreover, this property may lower the barrier to entry into omics research for non-specialists, particularly researchers with expertise in statistics or data science, and thereby facilitate more multidimensional development of omics studies. Furthermore, although the partitions identified by Monju were often interpretable in terms of cellular function, some partitions also spanned conventional cell-type boundaries. This feature may help move analyses beyond prior-knowledge-driven assumptions and toward data-structure-driven partitioning. In this sense, Monju may not only broaden access to omics research for non-specialists but also increase the flexibility of analyses conducted by domain experts.

A potential limitation of Monju is that its analysis explicitly depends on stochasticity across training runs. Although this stochasticity enables the quantification of cluster ambiguity and cellular-state continuity, it also raises the need to systematically evaluate stability and reproducibility. Future work should therefore examine how Monju’s run-to-run variability depends on dataset size, initialization, hyperparameters, and modality composition. An important direction for future work is to develop a single neural network framework that can directly learn multiple clustering criteria, without relying on repeated stochastic runs.

## 4 Methods

### 4.1 Monju model formulation

Monju is a deep generative mixture model that combines submodel assignment with submodel-specific variational autoencoding. For each cell, the model infers a discrete latent variable indicating the responsible submodel and a continuous latent variable within that submodel. Each submodel is implemented as a VAE and reconstructs multimodal count data through modality-specific decoders. The assignment network and all submodel-specific VAEs are trained jointly by maximizing an ELBO, so that cells that are well explained by a common low-dimensional generative structure tend to be assigned to the same submodel. For multimodal data, modality-specific encoders are combined using a mixture-of-experts-style inference model. We describe the model formally below.

#### 4.1.1 Notation

We consider multimodal single-cell count data *x* = (*x*_1_, *x*_2_, …, *x*_*M*_ ), where *M* denotes the number of modalities. The vector 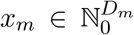 is the non-negative count vector for modality *m*, with dimensionality *D*_*m*_. In the CITE-seq analyses, *M* = 2, corresponding to RNA counts and Antibody-Derived Tags (ADT) counts.

Monju assumes that the cell population is generated from a mixture of *K* submodels. We introduce a discrete latent variable *k ∈* {1, …, *K*} that indicates the submodel responsible for given observation. For each submodel *k*, we introduce a continuous latent variable 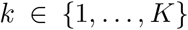, where *d*_*k*_ denotes the latent dimensionality of submodel *k*. We write the collection of all submodel-specific latent variables as *z*_1:*K*_ = (*z*_1_, *z*_2_, …, *z*_*K*_).

#### 4.1.2 Generative model

Monju is defined as a mixture of submodel-specific VAEs. Each submodel has its own latent variable and modality-specific decoders. For submodel *k*, the latent variable follows a diagonal Gaussian prior,

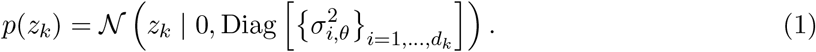

Given *z*_*k*_, each modality is generated independently through a modality-specific decoder,

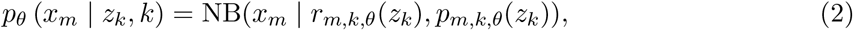

where NB indicates negative binomial distribution[11, 26]; *r*_*m,k,θ*_ and *p*_*m,k,θ*_ are decoder neural networks that output the parameters of a negative binomial distribution for modality *m*. The likelihood factorizes over modalities as

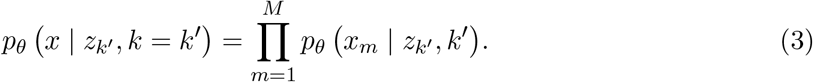

The submodel identity has a uniform prior,

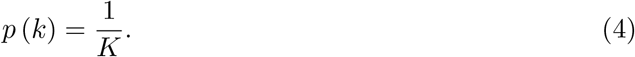

The marginal distribution of a cell is therefore a mixture of submodel-specific generative distributions,

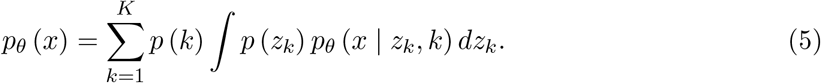

For variational inference, we write the joint distribution over all latent variables as

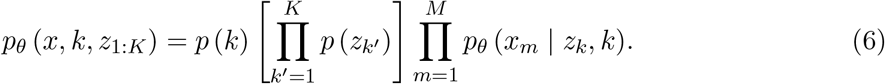

Only the latent variable corresponding to the selected submodel *k* contributes to generation of the observed cell. The remaining latent variables are auxiliary variables that follow their priors.

#### 4.1.3 Inference model

The exact posterior distribution

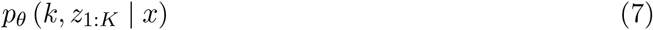

is intractable because the generative model uses neural decoders. We therefore use amortized variational inference. The variational distribution is defined as

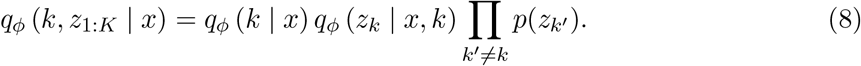

Thus, the inference model estimates the submodel assignment *k* and the latent variable *z*_*k*_ of the selected submodel. Latent variables associated with unselected submodels remain distributed according to their priors.

#### 4.1.4 Multimodal inference with mixture-of-experts encoders

For multimodal data, Monju uses modality-specific encoders. Each modality *m* produces both a submodel assignment distribution and submodel-specific Gaussian latent distributions:

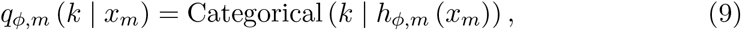

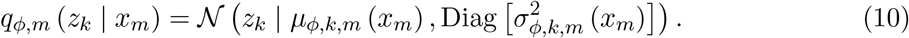

Here, *h*_*ϕ,m*_ maps modality *m* to a probability vector over submodels, and *µ*_*ϕ,k,m*_ and 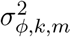 output the mean and diagonal variance of the latent distribution for submodel *k*, respectively. When multiple modalities are observed, Monju combines modality-specific inference distributions as uniformly weighted mixtures:

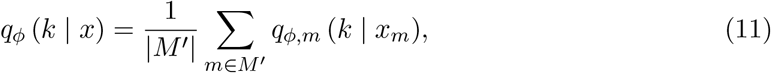

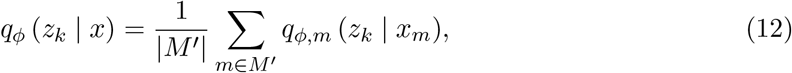

where *M* ^*′*^ *⊆* {1, …, *M* } is the set of observed modalities. This formulation also allows inference when some modalities are missing, by constructing the mixture only over available modalities.

#### 4.1.5 Variational objective

Monju is trained by maximizing the evidence lower bound. For a given submodel *k*, the submodel-specific ELBO for cell *x* is defined as

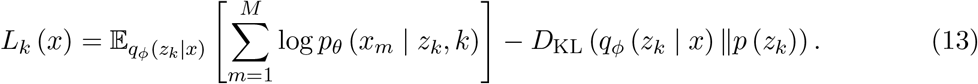

The full Monju objective marginalizes over submodel assignments:

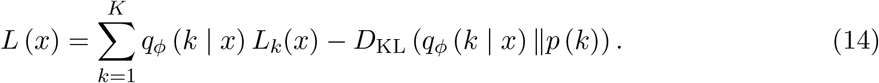

The first term evaluates how well each submodel reconstructs the cell, weighted by the inferred assignment probability. The second term regularizes the assignment distribution toward the prior over submodels. This objective is optimized jointly over the classifier network and all submodel-specific VAEs. The discrete expectation over *k* is computed exactly by summing over all submodels. The continuous latent expectation is estimated using Monte Carlo sampling with the reparameterization trick[27].

#### 4.1.6 Reconstruction and cross-modal prediction

We evaluated Monju by reconstruction and cross-modal prediction. Reconstruction refers to predicting an observed modality from the same input data. Cross-modal prediction refers to predicting an unobserved modality from other observed modalities. For example, in CITE-seq data, RNA can be used to predict surface-protein expression, or surface-protein expression can be used to predict RNA expression.

Given observed modalities *M* ^*′*^, we first select the most probable submodel,

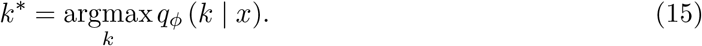

We then sample 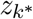 from the posterior distribution 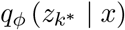 and generate predictions using the decoder of submodel *k*^*\**^. The mean of the negative binomial distribution is used as the point prediction.

#### 4.1.7 Meta-clustering across Monju runs

The assignment of cells to submodels is not uniquely determined due to inherent randomness in parameter initialization and stochastic sampling during training. As a result, Monju runs with different random seeds can produce different submodel partitions. We used this variability to integrate multiple clustering criteria.

We trained Monju models with 128 different random seeds. For each trial *t*, we obtained an assignment probability matrix

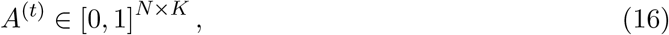

Where 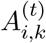 denotes the probability that cell *i* is assigned to submodel *k* in trial *t*.

For each cell *i* and trial *t*, we first defined its submodel assignment as a set, rather than as a single label, because some cells were assigned to multiple submodels with comparable probabilities. Specifically, we retained all submodels whose assignment probability was greater than or equal to 1*/*3:

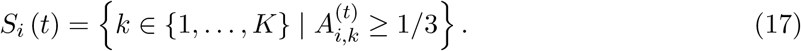

When two or more submodels were assigned with probabilities greater than or equal to 1*/*3, all of them were included in *S*_*i*_(*t*). For example, when *K* = 3, a cell with assignment probabilities [0.8, 0.1, 0.1] is represented as *S*_*i*_ (*t*) = *{*1*}*, whereas a cell with probabilities [0.4, 0.1, 0.5] is represented as *S*_*i*_ (*t*) = *{*1, 3*}*. Thus, *S*_*i*_(*t*) captures not only the most likely submodel but also ambiguity in submodel assignment when multiple submodels have similarly high probabilities.

For each cell, we concatenated these labels across trials:

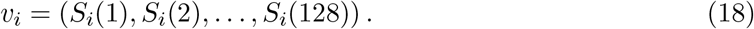

A meta-cluster was defined as a group of cells sharing the same label vector *v*_*i*_. Meta-clusters were ranked by cell number. For visualization and interpretation, meta-clusters containing fewer than 1,000 cells were grouped into the “Others” category.

#### 4.1.8 Quantification of meta-clustering ambiguity

To quantify ambiguity in meta-clustering, we evaluated how frequently each cell was assigned to the Others category. We first trained Monju with 128 different random seeds. We then repeatedly selected 12 distinct trials without replacement and performed meta-clustering using only those trials. In each meta-clustering run, meta-clusters containing fewer than 1,000 cells were labeled as Others. This procedure was repeated 100 times. For each cell *i*, we computed the fraction of runs in which the cell was assigned to Others.

#### 4.1.9 Construction and visualization of Monju Space

Monju Space was constructed from the classifier outputs across multiple Monju runs. For each cell and each trial, the classifier produced a *K*-dimensional assignment probability vector. We concatenated this *K*-dimensional vector across (*T* = 128) trials, yielding a (*K × T* )-dimensional representation:

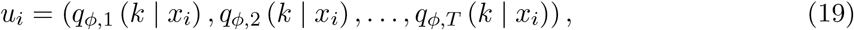

where *q*_*ϕ*_ (*k* | *x*_*i*_) is defined in Eq. (11). We defined this representation as Monju Space.

To visualize and characterize the structure of Monju Space, we applied UMAP to the Monju Space vectors and then performed Leiden clustering as a post hoc analysis. Leiden clustering in Figs. 5 and 6 was performed using the Python package `leidenalg` (version 0.10.1)[23]. For each input representation, we constructed an undirected weighted cell–cell graph using 30 nearest neighbors with Euclidean distance. The edge weight between two cells was defined as 1*/*(1 + *d*), where *d* denotes their Euclidean distance; when the same undirected edge appeared more than once, the larger weight was retained. Leiden partitions were obtained using `leidenalg.find_partition` with `RBConfigurationVertexPartition`, edge weights, and random seed 111. The resolution parameters were set to 0.5 and 1.0 for the reference latent-space analyses in Fig. 5c and Fig. 5d, respectively, and to 0.1 for the Monju Space analysis in Fig. 6b,d. Cells were visualized using both Leiden cluster labels and annotated cell-type labels. We also computed the composition of annotated cell types within each Leiden cluster and visualized the resulting matrix as a heatmap.

For comparison with a conventional clustering approach, we applied Leiden clustering to the reference latent space obtained from the conventional non-mixture VAE method using different resolution parameters. The resulting Leiden cluster labels were visualized on the same UMAP coordinates used for the Monju analyses. This comparison was used to evaluate how conventional distance-based clustering differs from clustering based on the partition-driven representation provided by Monju Space.

### 4.2 Dataset

We analyzed a peripheral blood mononuclear cell CITE-seq dataset from GEO accession GSE164378. We used RNA counts, Antibody-Derived Tags (ADT) counts, and metadata annotations. The raw dataset contained 161,764 cells, 33,538 RNA features, and 228 ADT features. After preprocessing, cells were split into training and held-out evaluation sets at an 8 : 2 ratio, resulting in 129,411 training cells and 32,353 evaluation cells.

### 4.3 Preprocessing

RNA count matrices were loaded into a Seurat object. Counts were log-normalized and highly variable genes (HVGs) were selected using the FindVariableFeatures function in Seurat v4[15, 28]. The top 5,000 HVGs were used for downstream analysis. Note that raw counts were used for Monju training. ADT counts were used without feature selection. No additional cell-level quality-control filtering was applied.

### 4.4 Network architecture

Each decoder consisted of one hidden block followed by two output heads. The hidden block was composed of a linear layer, batch normalization, and ReLU activation. One output head was clipped to the range [*−*12, 12] and then exponentiated to produce the negative binomial shape parameter *r*. The other output head was passed through a sigmoid function to produce the success-probability parameter *p. p* was clipped to the range [10^*−*8^, 1 *−* 10^*−*7^] .

The partitioning neural network for submodel assignment consisted of three hidden blocks, each composed of a linear layer, batch normalization, and ReLU activation, followed by an output layer. Outputs were clipped to the range [*−*12, 12] and then normalized using a softmax function to calculate the assignment probability *q* (*k* | *x*). For numerical stability, a small constant (10^*−*6^) was added to *q* (*k* | *x*).

The encoder networks for Gaussian latent variables also consisted of three hidden blocks followed by two output heads: one for the latent mean and one for the diagonal latent variance. For the variance, the output was clipped to the range [*−*12, 12], scaled by a softmax function, and then multiplied by the latent dimensionality. Latent variables were sampled by the reparameterization trick.

The main hyperparameters were as follows.

**Table 1.** Hyperparameters used in this study.

| Hyperparameter | Conventional VAE[12] | Monju |
| --- | --- | --- |
| Number of mixtures | 1 | 20 |
| Latent dimension | 10 | 10 |
| Hidden units | 200 | 200 |
| Decoder layers | 3 | 1 |
| Encoder layers | 3 | 3 |
| Batch size | 128 | 128 |
| Learning rate | $2 \times 10^{-3}$ | $2 \times 10^{-3}$ |
| Epochs | 50 | 50 |
| Warm-up | 25 | 25 |

For each modality *m*, we compute the library size

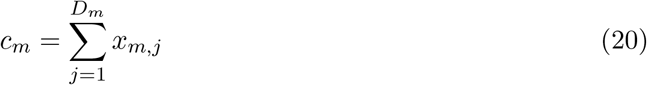

and apply normalization before passing the data into the encoder:

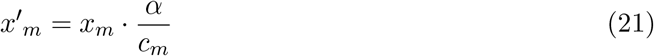

where *α* is a positive integer scale factor. On the decoder side, we restore the scale by rescaling the negative binomial parameter output by the decoder. In particular, letting *r* denote the decoder output corresponding to the NB shape/dispersion parameter, we compute:

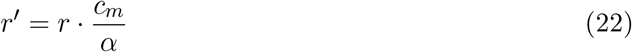

In this study, we assume that the target-modality library size *c*_*m*_ is given even in cross-modal prediction.

### 4.5 Model training

Monju was optimized using the Adam optimizer with AMSGrad[29]. The mini-batch size was 128, and the learning rate was 2 *×* 10^*−*3^. Training was performed for up to 50 epochs with early stopping based on the validation objective. The patience parameter for early stopping was 10 epochs.

To stabilize VAE training, deterministic warm-up[30] was used. The KL weight was linearly increased from 0 to 1 over the first 25 epochs. Gradients were clipped during training. Random seeds were fixed for training, data splitting, and visualization, although slight nondeterminism may remain due to GPU computation.

Unless otherwise stated, Monju was trained with (*K* = 20) submodels and latent dimensionality (*d*_*k*_ = 10) for each submodel. We trained Monju with 128 different random seeds to generate multiple partitions for meta-clustering.

### 4.6 Reference embedding

We used a 10-dimensional multimodal latent representation obtained from a conventional non-mixture VAE method (scMM)[12] as a reference representation. For visualization, UMAP[13] was applied to this representation to obtain two-dimensional coordinates.

### 4.7 of submodel assignments

To visualize Monju submodel assignments, we assigned each cell to the submodel with the maximum assignment probability and colored cells by this label on the reference UMAP. Submodels with no assigned cells after this maximum-probability assignment were excluded from visualization and composition summaries.

To quantify whether cells assigned to the same submodel were close in the reference latent space, we computed Euclidean distances between randomly sampled cell pairs within the same Monju submodel using the 10-dimensional reference latent representation. As a baseline, cell labels were randomly permuted while preserving submodel sizes, and the same distance calculation was repeated.

Cell-type composition within each Monju submodel was summarized by computing the fraction of each annotated cell type among cells assigned to each submodel.

#### 4.7.1 Mixed principal component (mPC)

To interpret latent structure within each submodel, we performed PCA on the submodel-specific latent representations. For each submodel, the first two principal component directions were denoted by *e*_1_ and *e*_2_.

We defined a one-dimensional mixed principal component axis as

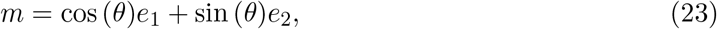

where *θ* is a submodel-specific rotation angle. The mPC score of cell *i* was defined as

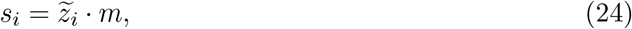

where 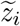 denotes the PCA-transformed coordinate of cell *i*.

The rotation angle was selected for each submodel by visual inspection of the PCA plot so that the axis captured the major biologically interpretable gradient within the submodel. For Fig. 4, we focused on submodel *z*_2_, which mainly contained CD4^+^ T cells.

#### 4.7.2 Molecular variation along the mPC axis

We characterized molecular variation along the mPC axis from two complementary perspectives: (i) expression decoded from the trained generative model, and (ii) raw observed expression. The former reveals what the model has learned to encode along the axis, whereas the latter verifies that the same gradient is present in the data without invoking the decoder.

To examine how gene and protein expression varied along the mPC axis, positions were sampled uniformly along the axis. For each sampled position, a latent vector was constructed and decoded using the trained decoder of the corresponding submodel.

Reconstructed RNA and protein expression values were log1p-transformed and centered across sampled positions. Molecules were ranked by the covariance between reconstructed expression and mPC position; the top 12 with positive and top 12 with negative covariance were visualized as heatmaps. The selected molecules were visualized as heatmaps.

Observed expression patterns along the mPC axis were also evaluated. For each gene and protein, the Spearman rank correlation coefficient (*ρ*) between observed log1p-transformed counts and mPC scores was computed. Among the features with a false discovery rate below 0.05, the top 40 genes and top 40 proteins ranked by the absolute value of *ρ* were visualized.

## Supplementary information

Supplementary information is available for this paper and includes Supplementary Figs. S1–S16 and additional analyses supporting the main conclusions. The supplementary material contains evaluations of cross-modal prediction tasks, verification of submodel assignments, comparisons with data labels, correlations between mPC scores and observed expression, and stability analyses across random seeds.

## Acknowledgements

This study was supported in part by the Moonshot R&D Program (JPMJMS2024-9 to H.N.), CREST (JPMJCR25Q2 to H.N.), and the Establishment of University Fellowships Towards the Creation of Science, Technology and Innovation (JPMJFS2129 to T.K.) from the Japan Science and Technology Agency (JST), as well as Multidisciplinary Frontier Brain and Neuroscience Discoveries (Brain/MINDS 2.0) (JP25wm0625322 and JP25wm0625210 to H.N.) from the Japan Agency for Medical Research and Development (AMED).

## Declarations

### Competing interests

The authors declare no competing interests.

### Data availability

This study analyzed a previously published, publicly available PBMC CITE-seq dataset from GEO under accession GSE164378. No new sequencing data were generated in this study. Processed data generated during this study are available from the corresponding authors upon reasonable request.

### Code availability

The Python (v3.9.7) code used to train Monju and generate the figures will be made available on GitHub upon publication.

### Author contributions

T.K., S.S., and H.N. conceived the study. T.K., S.S., and H.N. developed the method. T.K. performed the computational analyses with assistance from S.S., and R.K., S.F., and Y.Y. contributed to data interpretation. T.K., S.S., and H.N. drafted the manuscript. All authors discussed the results, revised the manuscript, and approved the final version.

## Appendix A Results on cross-modal prediction tasks

**Figure S1.**
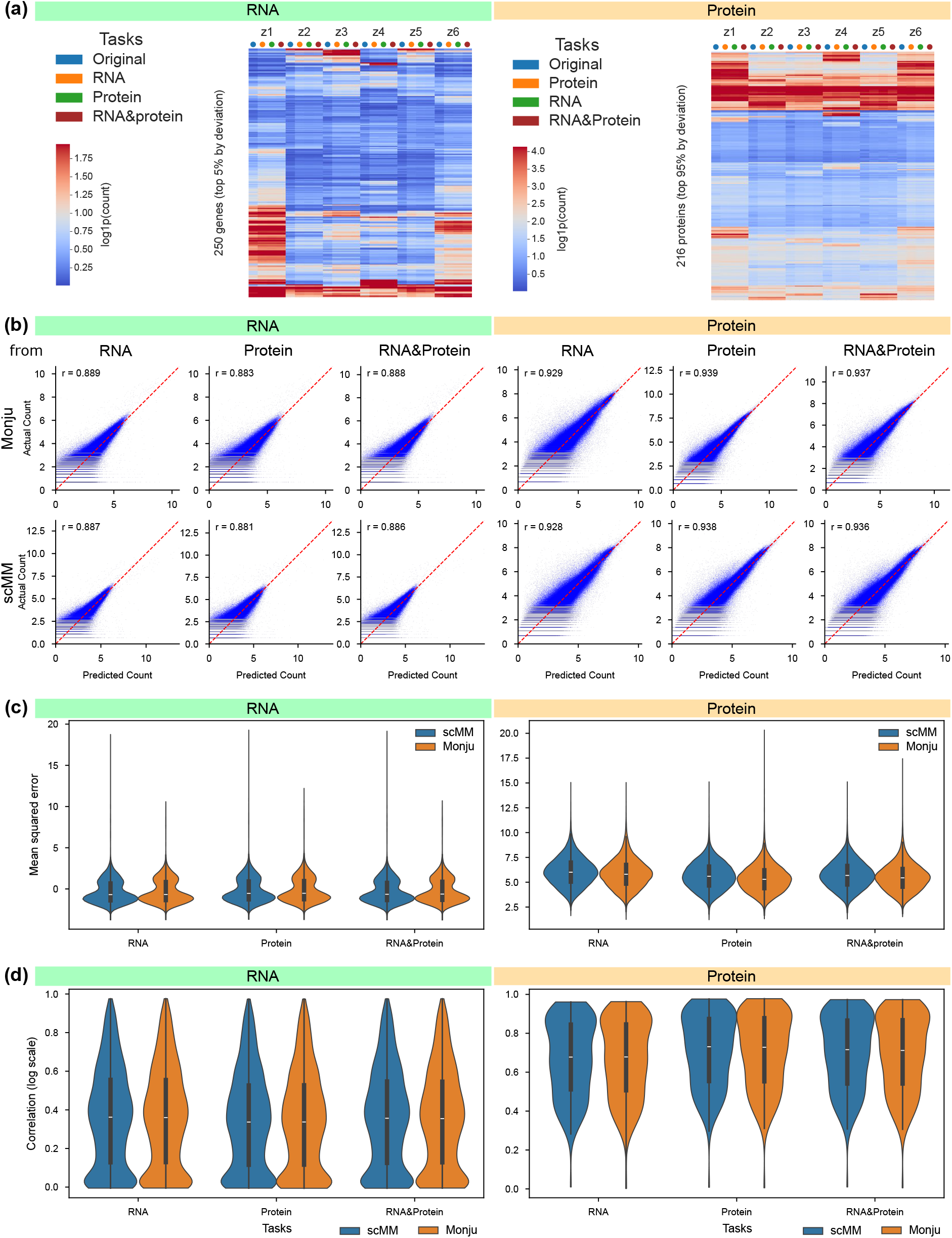
Validation of cross-modal prediction error. **(a)** Mean log-scale RNA/protein expression per submodel across original data, single-modality reconstruction, cross-modal prediction, and reconstruction from both modalities. **(b)** Reconstructed/predicted versus observed log-scale expression for RNA and protein. Monju and scMM are shown in the top and bottom rows, respectively; Pearson correlation coefficients are annotated in each panel. **(c)** Comparison of RNA and protein prediction errors between scMM and Monju. **(d)** Distributions of per-RNA/per-protein Pearson correlation coefficients for each prediction task.

## Appendix B Verification of submodel assignments by Monju

**Figure S2.**
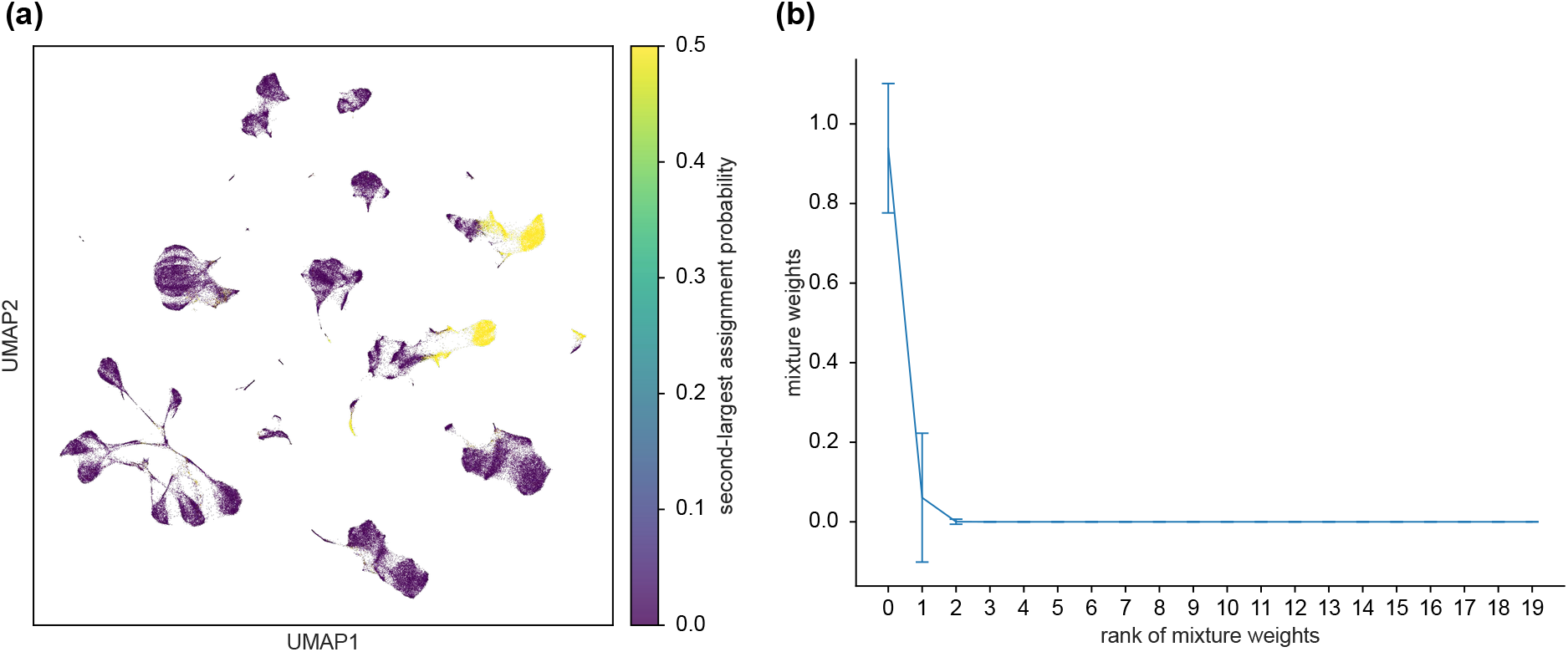
Distribution of Monju submodel assignments. **(a)** The second-largest assignment probability over Monju’s submodels shown on the same UMAP plots as Fig. 3a. **(b)** Mean assignment probability versus rank: for each cell, assignment probabilities across submodels are sorted in descending order; values are then averaged across all cells at each rank. The horizontal axis is the rank of the assignment probability and the vertical axis is the cell-averaged probability.

## Appendix C Comparison of submodel assignments with data labels

**Figure S3.**
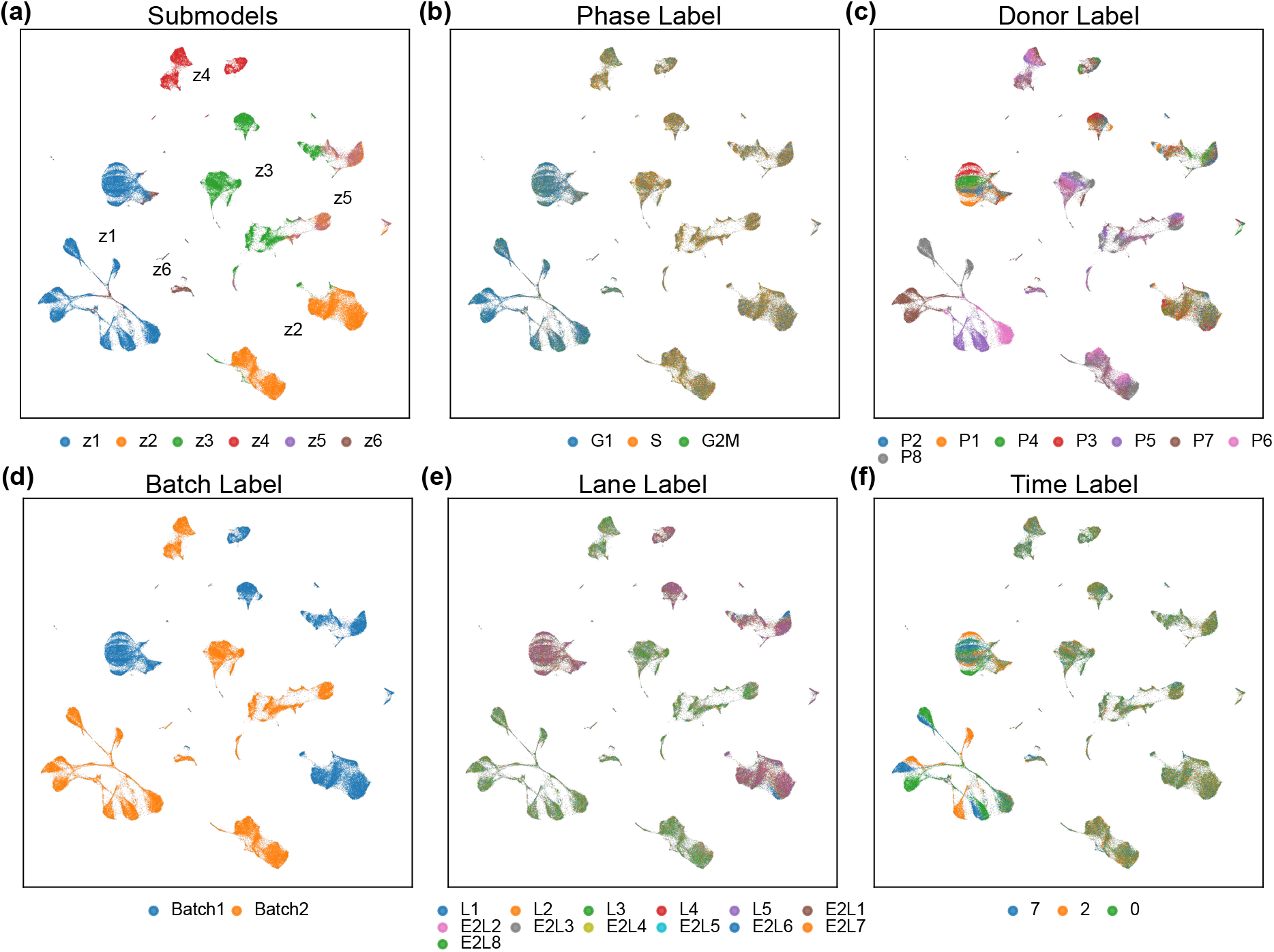
Visualization of meta-data labels. (a) Reproduction of Fig. 3a showing Monju submodel assignments. (b-f) Metadata labels for the PBMC scCITE-seq dataset visualized on the same UMAP embedding as in Fig. 3a. Points are colored according to (b) cell-cycle phase, (c) Donor ID, (d) Batch ID, (e) 10x Chromium chip lane ID, and (f) days since vaccination (time labels). In each panel, the legend lists the corresponding label categories and the number of cells assigned to each category.

**Figure S4.**
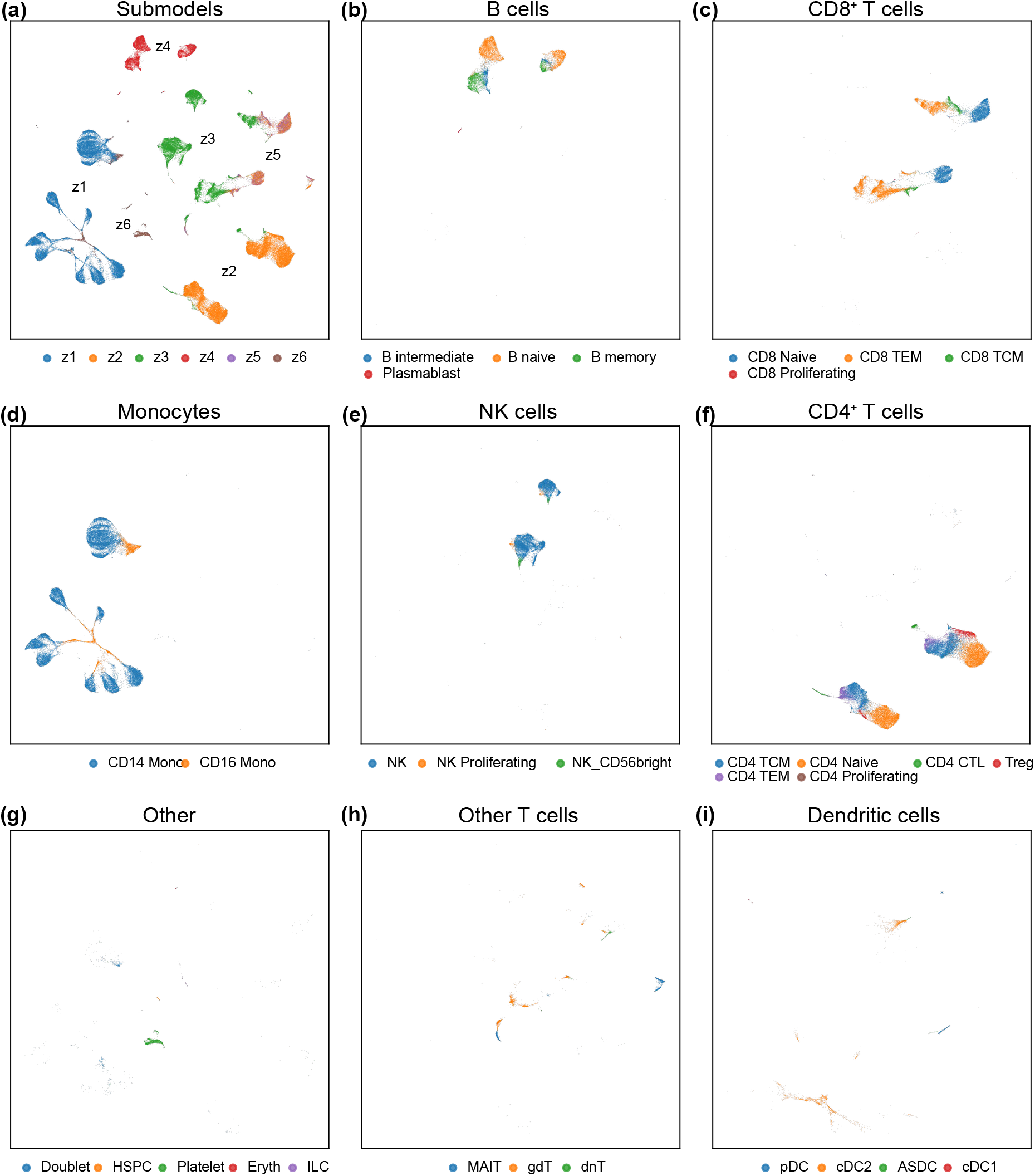
Visualization of cell-type labels. (a) Reproduction of Fig. 3a showing Monju submodel assignments. (b-i) Fine-grained cell-type labels (celltype.l2) of the PBMC scCITE-seq dataset visualized on the same UMAP embedding as in Fig. 3a. Cells were plotted separately for each celltype.l1 label, and points are colored by celltype.l2. The upper-left legend lists the celltype.l2 categories and the number of cells assigned to each category. Panels show cells annotated as (b) B cells, (c) CD8 T cells, (d) Monocytes, (e) NK cells, (f) CD4 T cells, (g) other, (h) other T cells, and (i) DC cells.

## Appendix D Analysis of latent spaces for submodels other than

**Figure S5.**
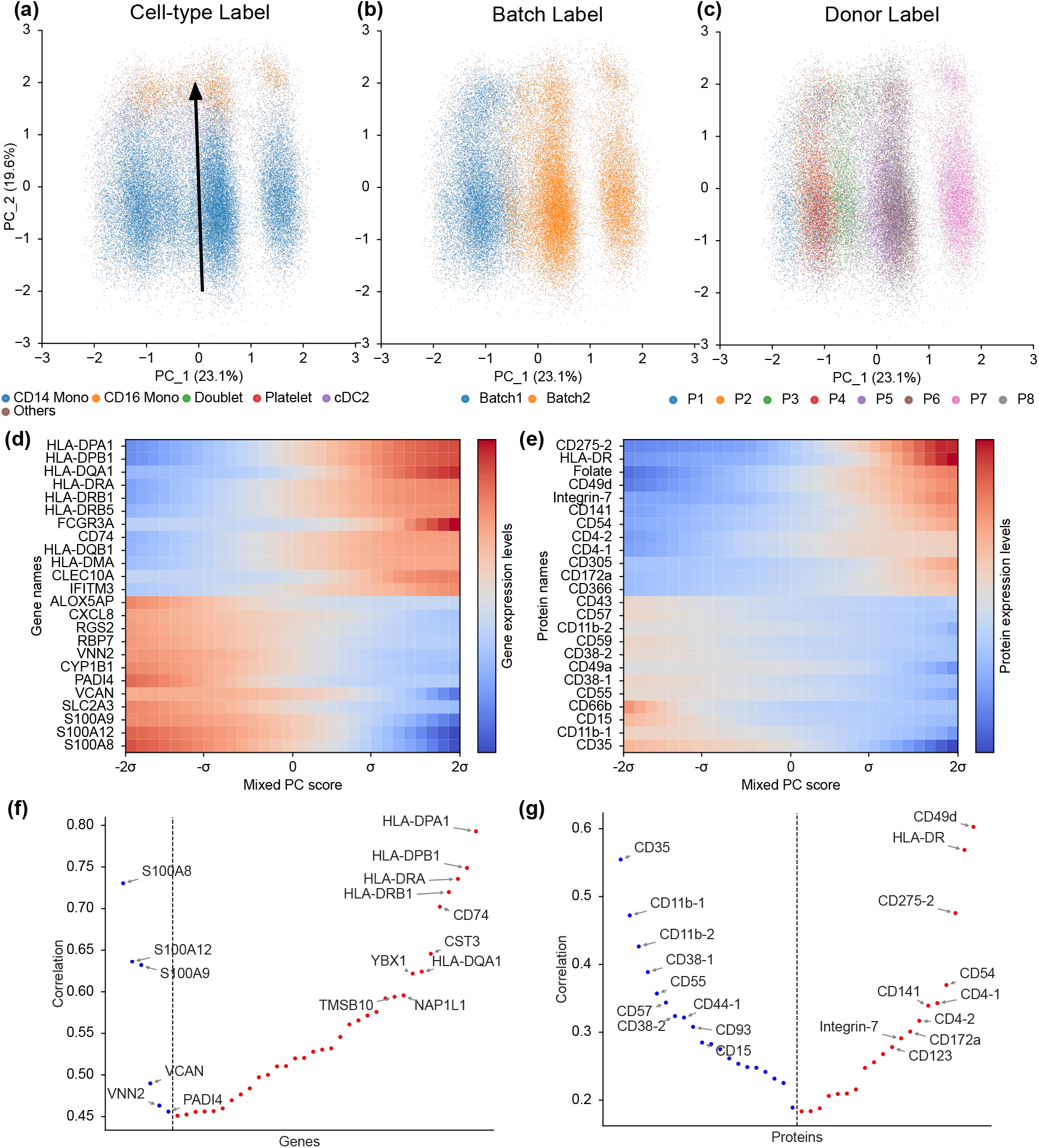
Interpretability of the latent representation of *z*_1_ learned by Monju. **(a,b, and c)** PCA visualization of submodel *z*_1_. Only cells assigned to *z*_1_ were extracted, and PCA was applied to their latent representations within this submodel. Points represent individual cells and are colored by their **(a)** cell-type labels, batch ID, and **(c)** donor ID, respectively. The arrow in (a) shows the axis used for visualizing expression changes in panel (d and e). **(d and e)** Heatmaps showing gene **(d)** and protein **(e)** expression changes along the axis indicated in (a). The horizontal axis represents positions along the axis. Expression values at each position were estimated using the corresponding submodel VAE by decoding latent coordinates sampled along this axis. The vertical axis lists genes or proteins. Labels shown in blue indicate molecules that also appear in (f and g). Expression values are shown on a log scale and normalized to have a mean of zero for each gene or protein. **(f and g)** Genes **(f)** and proteins **(g)** whose observed expression levels correlate with the mixed principal component in (a). Genes and proteins are ordered along the horizontal axis by ascending Spearman correlation coefficient. The vertical axis indicates the absolute Spearman correlation between the mPC score and observed expression. Positive correlations are shown in red and negative correlations in blue. Only the top 40 genes and top 40 proteins with the largest absolute correlations are displayed.

**Figure S6.**
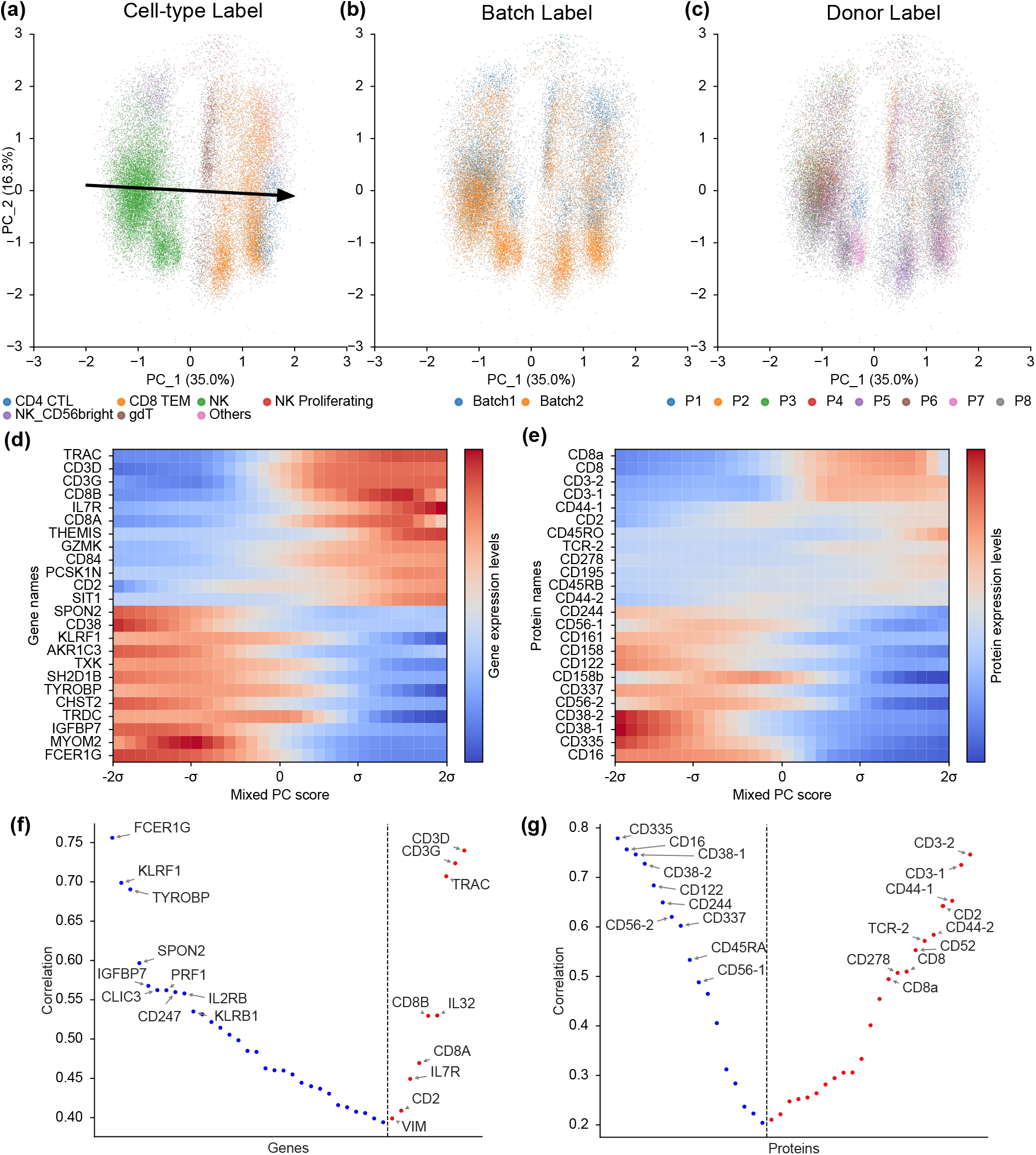
Interpretability of the latent representation of *z*_3_ learned by Monju. **(a,b, and c)** PCA visualization of submodel *z*_3_. Only cells assigned to *z*_3_ were extracted, and PCA was applied to their latent representations within this submodel. Points represent individual cells and are colored by their **(a)** cell-type labels, **(b)** batch ID, and **(c)** donor ID, respectively. The arrow in (a) shows the axis used for visualizing expression changes in panel (d and e). **(d and e)** Heatmaps showing gene **(d)** and protein **(e)** expression changes along the axis indicated in (a). The horizontal axis represents positions along the axis. Expression values at each position were estimated using the corresponding submodel VAE by decoding latent coordinates sampled along this axis. The vertical axis lists genes or proteins. Labels shown in blue indicate molecules that also appear in (f and g). Expression values are shown on a log scale and normalized to have a mean of zero for each gene or protein. **(f and g)** Genes **(f)** and proteins **(g)** whose observed expression levels correlate with the mixed principal component in (a). Genes and proteins are ordered along the horizontal axis by ascending Spearman correlation coefficient. The vertical axis indicates the absolute Spearman correlation between the mPC score and observed expression. Positive correlations are shown in red and negative correlations in blue. Only the top 40 genes and top 40 proteins with the largest absolute correlations are displayed.

**Figure S7.**
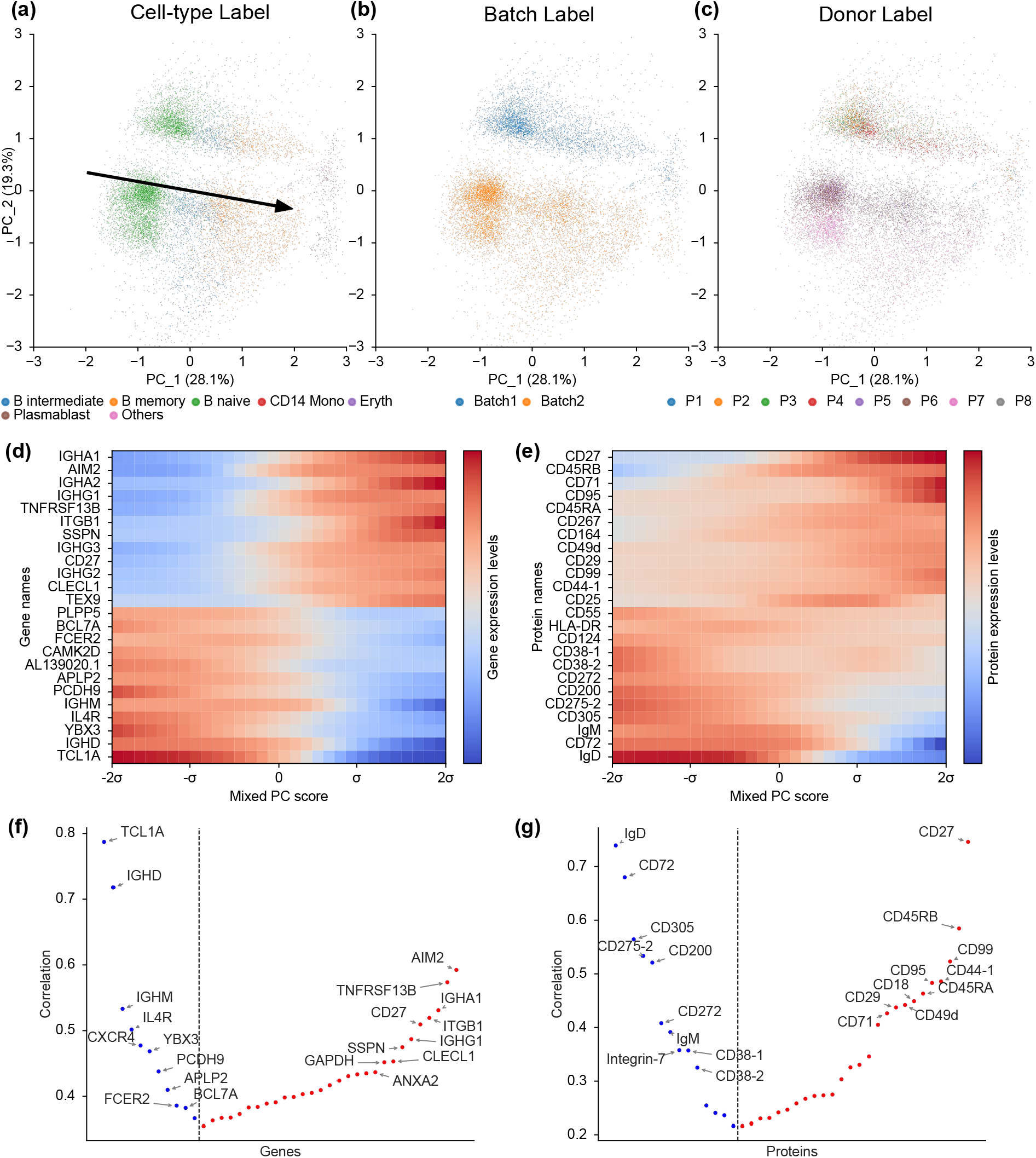
Interpretability of the latent representation of *z*_4_ learned by Monju. **(a,b, and c)** PCA visualization of submodel *z*_4_. Only cells assigned to *z*_4_ were extracted, and PCA was applied to their latent representations within this submodel. Points represent individual cells and are colored by their **(a)** cell-type labels, **(b)** batch ID, and **(c)** donor ID, respectively. The arrow in (a) shows the axis used for visualizing expression changes in panel (d and e). **(d and e)** Heatmaps showing gene **(d)** and protein **(e)** expression changes along the axis indicated in (a). The horizontal axis represents positions along the axis. Expression values at each position were estimated using the corresponding submodel VAE by decoding latent coordinates sampled along this axis. The vertical axis lists genes or proteins. Labels shown in blue indicate molecules that also appear in (f and g). Expression values are shown on a log scale and normalized to have a mean of zero for each gene or protein. **(f and g)** Genes **(f)** and proteins **(g)** whose observed expression levels correlate with the mixed principal component in (a). Genes and proteins are ordered along the horizontal axis by ascending Spearman correlation coefficient. The vertical axis indicates the absolute Spearman correlation between the mPC score and observed expression. Positive correlations are shown in red and negative correlations in blue. Only the top 40 genes and top 40 proteins with the largest absolute correlations are displayed.

**Figure S8.**
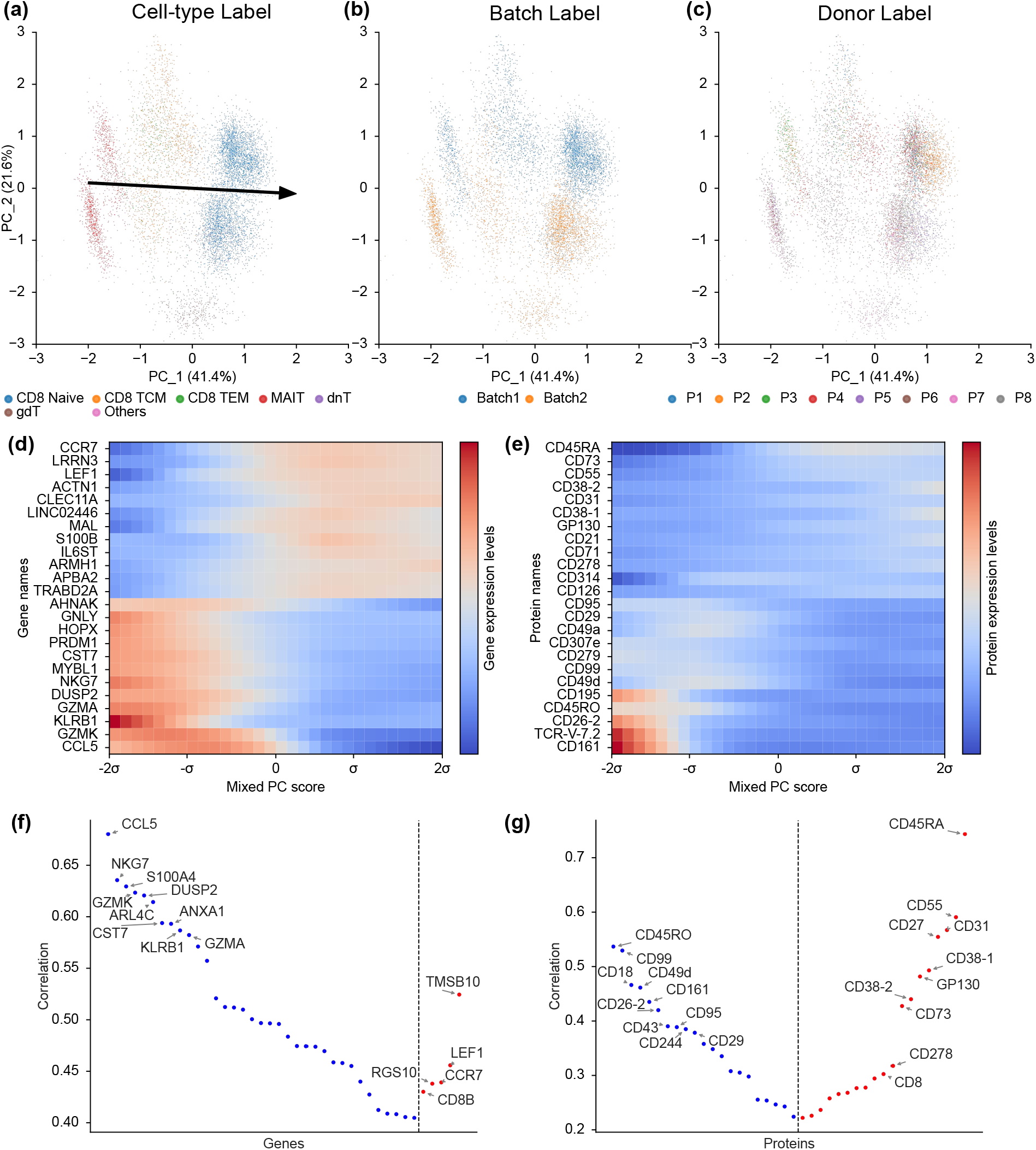
Interpretability of the latent representation of *z*_5_ learned by Monju. **(a,b, and c)** PCA visualization of submodel *z*_5_. Only cells assigned to *z*_5_ were extracted, and PCA was applied to their latent representations within this submodel. Points represent individual cells and are colored by their **(a)** cell-type labels, **(b)** batch ID, and **(c)** donor ID, respectively. The arrow in (a) shows the axis used for visualizing expression changes in panel (d and e). **(d and e)** Heatmaps showing gene **(d)** and protein **(e)** expression changes along the axis indicated in (a). The horizontal axis represents positions along the axis. Expression values at each position were estimated using the corresponding submodel VAE by decoding latent coordinates sampled along this axis. The vertical axis lists genes or proteins. Labels shown in blue indicate molecules that also appear in (f and g). Expression values are shown on a log scale and normalized to have a mean of zero for each gene or protein. **(f and g)** Genes **(f)** and proteins **(g)** whose observed expression levels correlate with the mixed principal component in (a). Genes and proteins are ordered along the horizontal axis by ascending Spearman correlation coefficient. The vertical axis indicates the absolute Spearman correlation between the mPC score and observed expression. Positive correlations are shown in red and negative correlations in blue. Only the top 40 genes and top 40 proteins with the largest absolute correlations are displayed.

**Figure S9.**
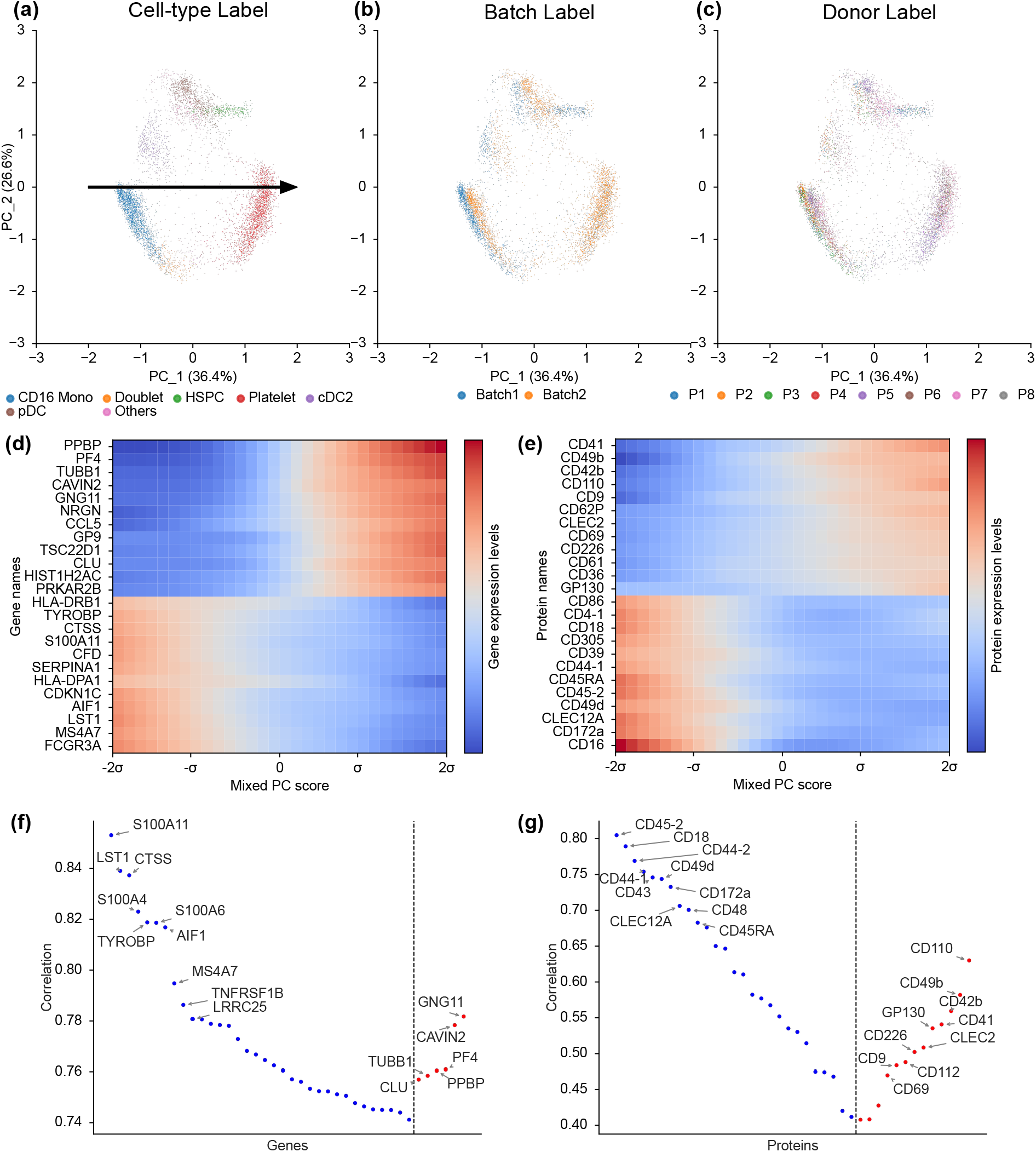
Interpretability of the latent representation of *z*_6_ learned by Monju. **(a,b, and c)** PCA visualization of submodel *z*_6_. Only cells assigned to *z*_6_ were extracted, and PCA was applied to their latent representations within this submodel. Points represent individual cells and are colored by their **(a)** cell-type labels, **(b)** batch ID, and **(c)** donor ID, respectively. The arrow in (a) shows the axis used for visualizing expression changes in panel (d and e). **(d and e)** Heatmaps showing gene **(d)** and protein **(e)** expression changes along the axis indicated in (a). The horizontal axis represents positions along the axis. Expression values at each position were estimated using the corresponding submodel VAE by decoding latent coordinates sampled along this axis. The vertical axis lists genes or proteins. Labels shown in blue indicate molecules that also appear in (f and g). Expression values are shown on a log scale and normalized to have a mean of zero for each gene or protein. **(f and g)** Genes **(f)** and proteins **(g)** whose observed expression levels correlate with the mixed principal component in (a). Genes and proteins are ordered along the horizontal axis by ascending Spearman correlation coefficient. The vertical axis indicates the absolute Spearman correlation between the mPC score and observed expression. Positive correlations are shown in red and negative correlations in blue. Only the top 40 genes and top 40 proteins with the largest absolute correlations are displayed.

## Appendix E Relationship between submodels and data labels

**Figure S10.**
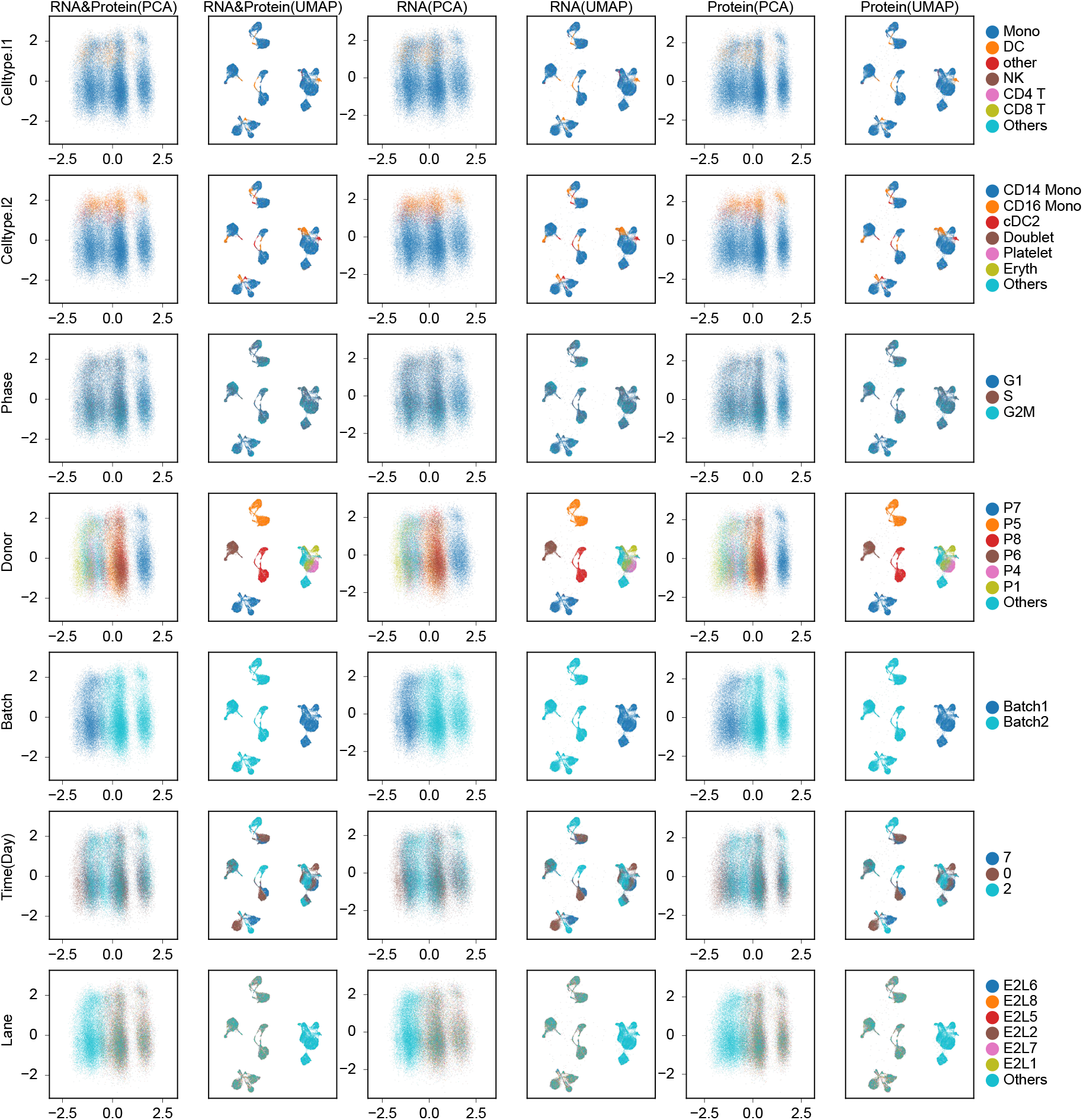
Visualization of submodel 1 latent spaces colored by each metadata label. Cells assigned to submodel *z*_1_ were extracted, and their submodel-specific latent representations were computed using the modality information indicated for each column. The six columns show PCA or UMAP projections based on RNA and protein information, RNA information alone, or protein information alone. From left to right, the columns correspond to RNA & Protein-PCA, RNA & Protein-UMAP, RNA-PCA, RNA-UMAP, protein-PCA, and protein-UMAP. Each row shows the same cells colored by a different data label: celltype.l1, celltype.l2, cell-cycle phase, donor ID, batch ID, days since vaccination, and 10x Chromium chip lane ID. The legend on the right lists the label categories and the number of cells assigned to each category.

**Figure S11.**
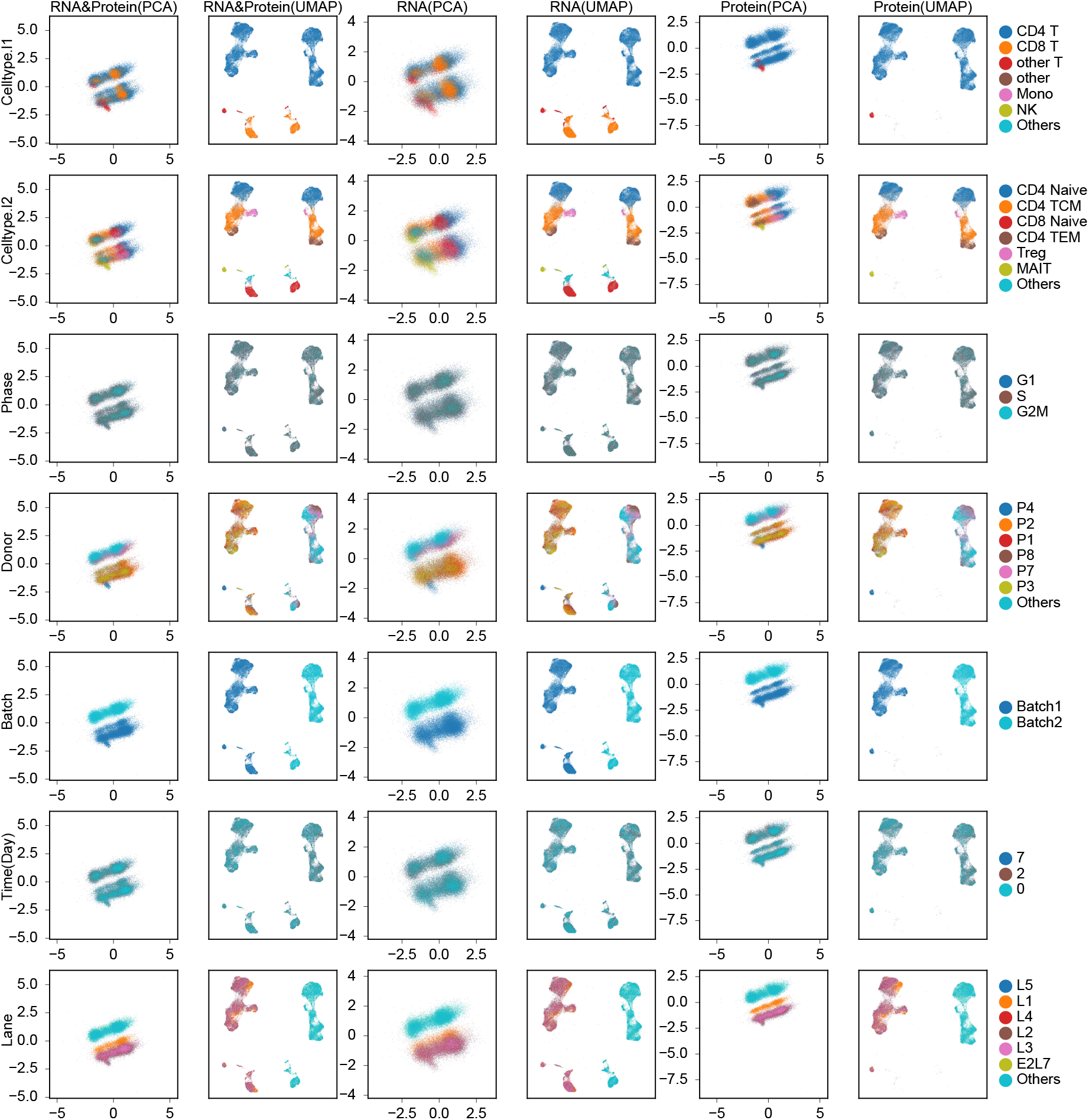
Visualization of submodel 2 latent spaces colored by each metadata label. Cells assigned to submodel *z*_2_ were extracted, and their submodel-specific latent representations were computed using the modality information indicated for each column. The six columns show PCA or UMAP projections based on RNA and protein information, RNA information alone, or protein information alone. From left to right, the columns correspond to RNA & Protein-PCA, RNA & Protein-UMAP, RNA-PCA, RNA-UMAP, protein-PCA, and protein-UMAP. Each row shows the same cells colored by a different data label: celltype.l1, celltype.l2, cell-cycle phase, donor ID, batch ID, days since vaccination, and 10x Chromium chip lane ID. The legend on the right lists the label categories and the number of cells assigned to each category.

**Figure S12.**
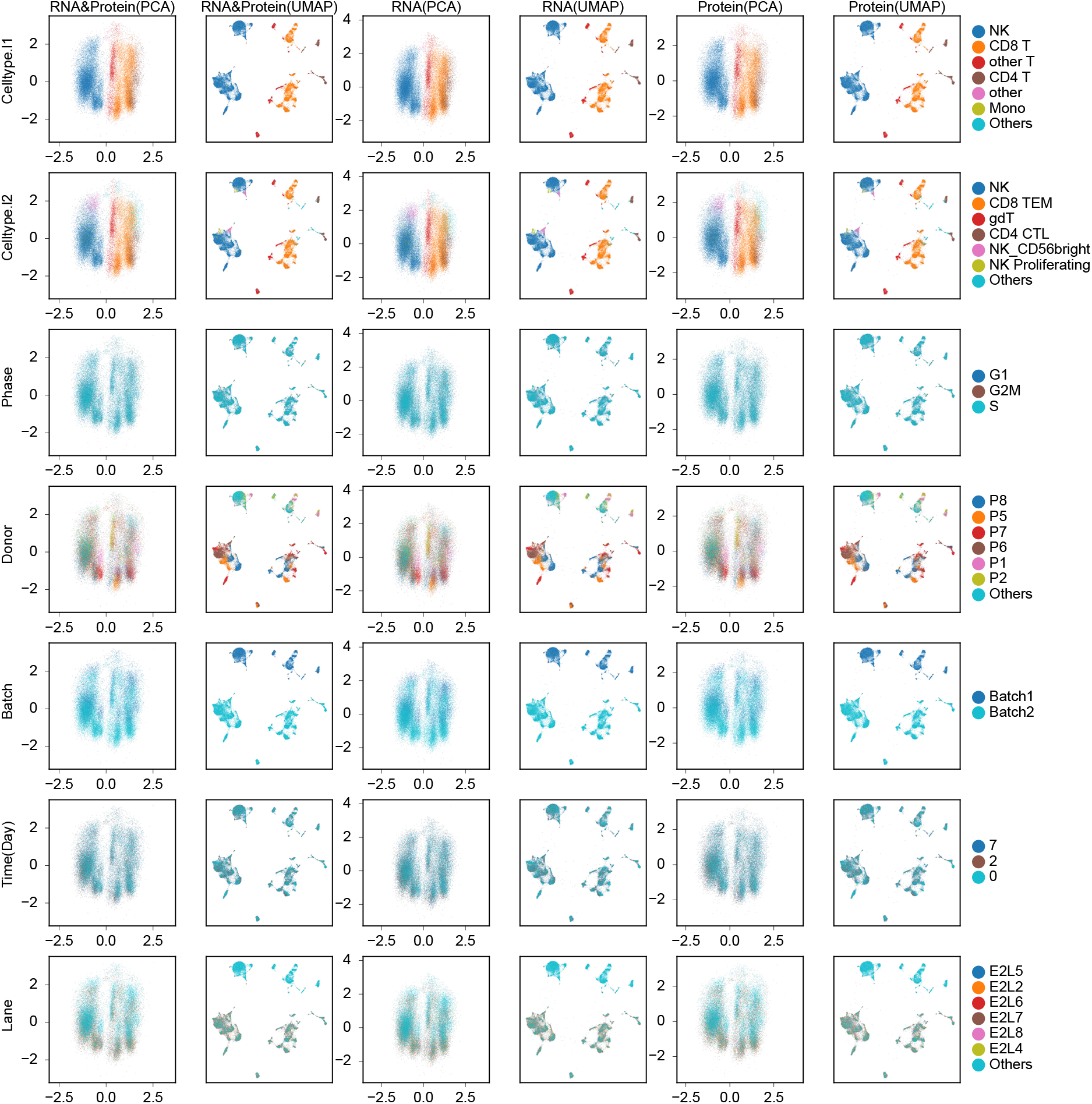
Visualization of submodel 3 latent spaces colored by each metadata label. Cells assigned to submodel *z*_3_ were extracted, and their submodel-specific latent representations were computed using the modality information indicated for each column. The six columns show PCA or UMAP projections based on RNA and protein information, RNA information alone, or protein information alone. From left to right, the columns correspond to RNA & Protein-PCA, RNA & Protein-UMAP, RNA-PCA, RNA-UMAP, protein-PCA, and protein-UMAP. Each row shows the same cells colored by a different data label: celltype.l1, celltype.l2, cell-cycle phase, donor ID, batch ID, days since vaccination, and 10x Chromium chip lane ID. The legend on the right lists the label categories and the number of cells assigned to each category.

**Figure S13.**
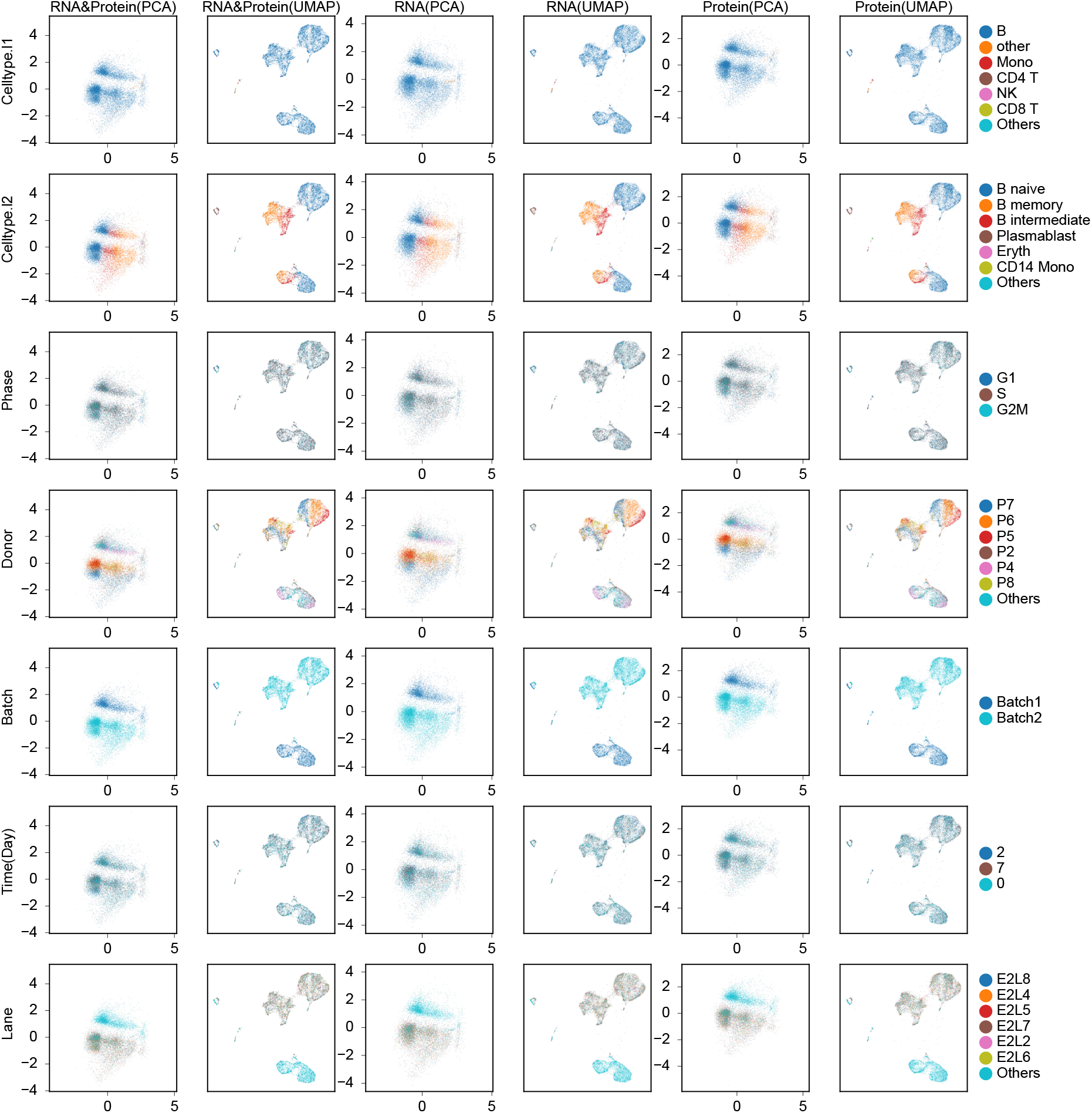
Visualization of submodel 4 latent spaces colored by each metadata label. Cells assigned to submodel *z*_4_ were extracted, and their submodel-specific latent representations were computed using the modality information indicated for each column. The six columns show PCA or UMAP projections based on RNA and protein information, RNA information alone, or protein information alone. From left to right, the columns correspond to RNA & Protein-PCA, RNA & Protein-UMAP, RNA-PCA, RNA-UMAP, protein-PCA, and protein-UMAP. Each row shows the same cells colored by a different data label: celltype.l1, celltype.l2, cell-cycle phase, donor ID, batch ID, days since vaccination, and 10x Chromium chip lane ID. The legend on the right lists the label categories and the number of cells assigned to each category.

**Figure S14.**
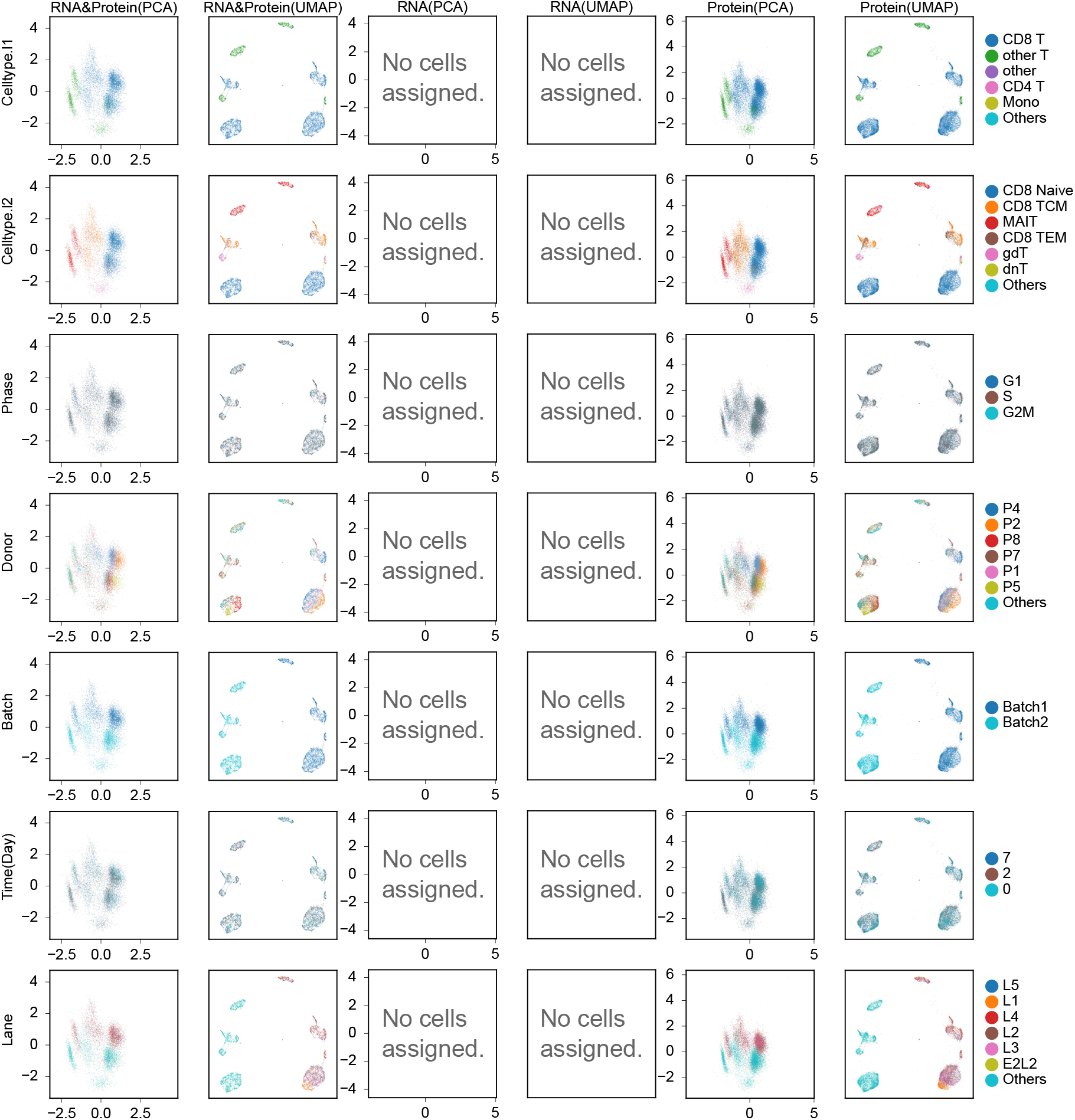
Visualization of submodel 5 latent spaces colored by each metadata label. Cells assigned to submodel *z*_5_ were extracted, and their submodel-specific latent representations were computed using the modality information indicated for each column. The six columns show PCA or UMAP projections based on RNA and protein information, RNA information alone, or protein information alone. From left to right, the columns correspond to RNA & Protein-PCA, RNA & Protein-UMAP, RNA-PCA, RNA-UMAP, protein-PCA, and protein-UMAP. Each row shows the same cells colored by a different data label: celltype.l1, celltype.l2, cell-cycle phase, donor ID, batch ID, days since vaccination, and 10x Chromium chip lane ID. The legend on the right lists the label categories and the number of cells assigned to each category.

**Figure S15.**
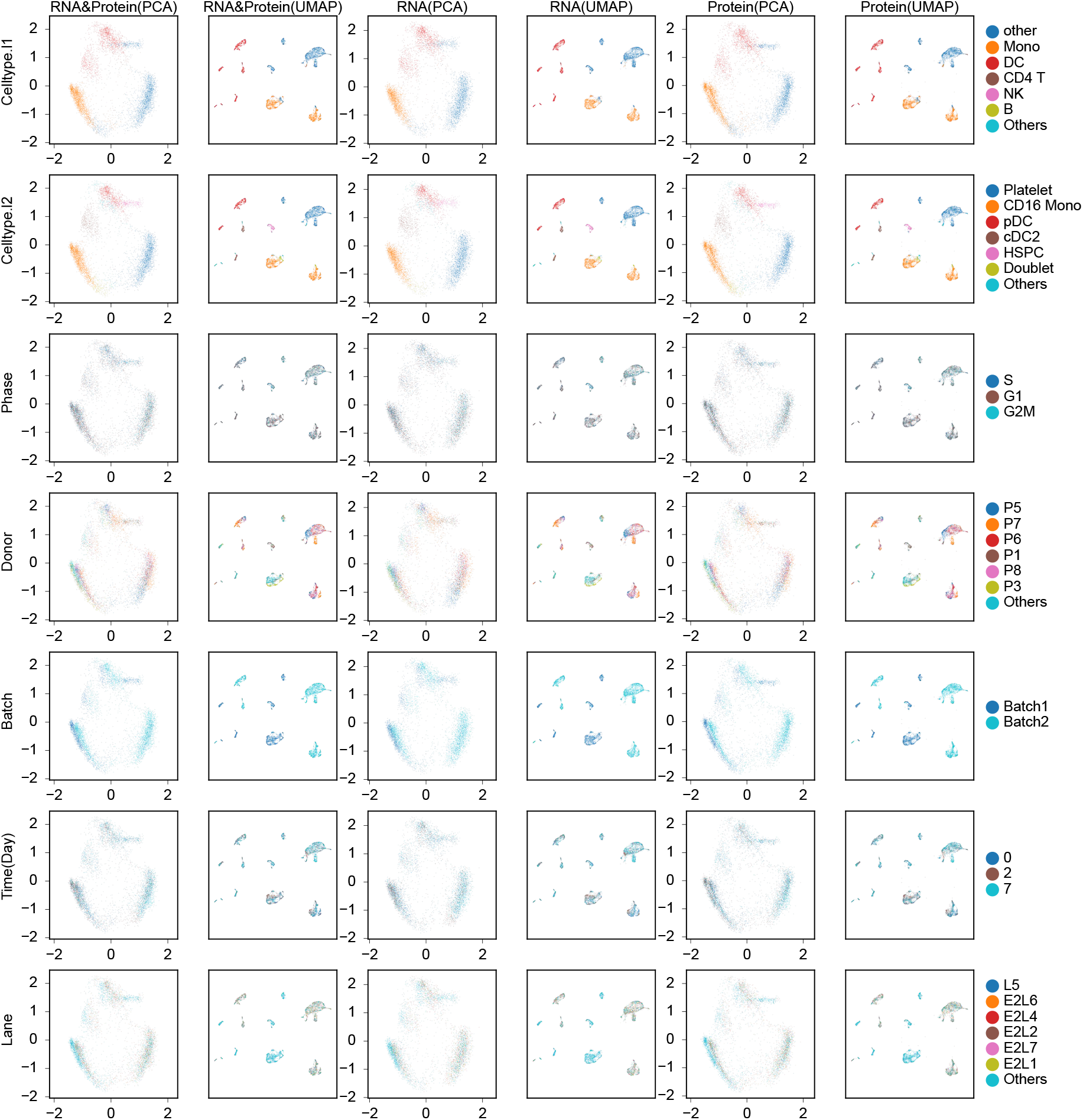
Visualization of submodel 6 latent spaces colored by each metadata label. Cells assigned to submodel *z*_6_ were extracted, and their submodel-specific latent representations were computed using the modality information indicated for each column. The six columns show PCA or UMAP projections based on RNA and protein information, RNA information alone, or protein information alone. From left to right, the columns correspond to RNA & protein-PCA, RNA & protein-UMAP, RNA-PCA, RNA-UMAP, protein-PCA, and protein-UMAP. Each row shows the same cells colored by a different data label: celltype.l1, celltype.l2, cell-cycle phase, donor ID, batch ID, days since vaccination, and 10x Chromium chip lane ID. The legend on the right lists the label categories and the number of cells assigned to each category.

## APPENDIX F Stability of Monju submodel assignments

**Figure S16.**
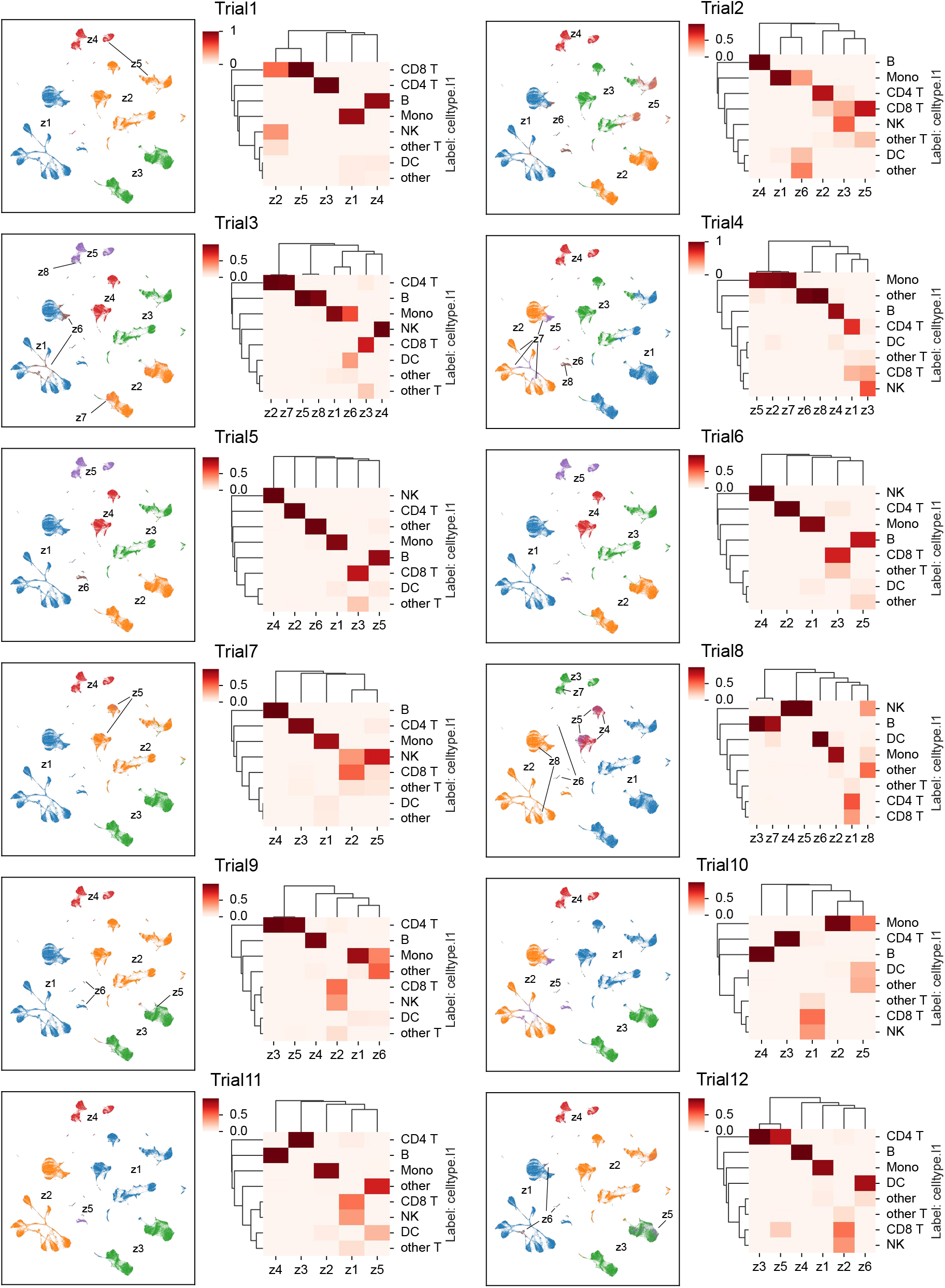
Stability of submodel assignments produced by Monju. Monju learning results across trials with different random seeds, shown as pairs of panels per trial. Left panel: Submodel IDs assigned by Monju, visualized on the same UMAP plots as Fig. 3a. Color indicates the submodel ID assigned by Monju. Right panel: Heatmap showing, for each submodel, the proportion of cells of each cell type. The vertical axis lists cell-type labels (celltype.l1) and the horizontal axis lists submodel IDs. Cell types with higher proportions are shown in darker shades, as indicated by the color bar.

## Notes

### Competing Interest Statement

The authors have declared no competing interest.

